# Distinct region-specific neutralization profiles of contemporary HIV-1 clade C against best-in-class broadly neutralizing antibodies

**DOI:** 10.1101/2024.12.31.630867

**Authors:** Jyoti Sutar, Priyanka Jayal, Ranajoy Mullick, Sangeeta Chaudhary, Prajakta Kamble, Shilpa Bhowmick, Snehal Kaginkar, Varsha Padwal, Pratik Devadiga, Namrata Neman, Dale Kitchin, Haajira Kaldine, Nonhlanhla N. Mkhize, Bongiwe Ndlovu, Kamini Gounder, Sohini Mukherjee, Shweta Shrivas, Neha Sharma, Chaman Prasad, Sonia Tewatia, Nainika Parihar, Naresh Kumar, Nandini Kasarpalkar, Balwant Singh, Shobha Mohapatra, Mohammad Aquil, C. Vishal Kumar, Thongadi Ramesh Dinesha, Aylur Kailasom Srikrishnan, Jayanthi Shastri, Sachee Agrawal, Sushma Gaikwad, Sayantani Mondal, Bhaswati Bandopadhyay, Subhasish Kamal Guha, Dipesh Kale, Debashis Biswas, Dhanashree Patil, Ramesh S. Paranjape, Satyajit Mukhopadhyay, Hema, Ritika Das, Anand Kondapi, Vikrant Bhor, Suprit Deshpande, Devin Sok, Thumbi Ndung’u, Penny L Moore, Kailapuri Gangatharan Murugavel, Vainav Patel, Jayanta Bhattacharya

## Abstract

Broadly neutralizing antibodies (bnAb) have been clinically proven to be an excellent choice for HIV-1 prevention. However, the relative effectiveness of best-in-class bnAbs against regionally relevant circulating HIV-1 forms is not clear. In the present study, we compared the degree of neutralization sensitivity of contemporary HIV-1 Indian clade C with that of South African origin. Phylogenetic analysis revealed that these clade C viruses continue to evolve distinctly from one another. Env-pseudotyped viruses prepared using contemporary HIV-1 clade C *env* genes (N=115) obtained from nine geographically distinct sites in India (between 2020-2023) were found to be most sensitive to V3-directed bnAbs 10-1074 and BG18, and second generation CD4 binding site (CD4bs) directed bnAbs (VRC07, N6 and 1-18), however they were found to be significantly resistant to V1/V2 apex directed bnAbs. Moreover, we observed that the degree of sensitivity varied between contemporary Indian and South African clade C viruses. Differences in degree of neutralization susceptibility were associated with differences observed in key residues that form bnAb contact sites, gp120 loop lengths and the number of N-linked glycans in the V4 hypervariable region. Interestingly, the second generation CD4bs bnAbs (VRC07, N6, 1-18) showed neutralization of VRC01 and 3BNC117 resistant viruses but with 2-7-fold reduced potency compared to the VRC01 sensitive counterparts, likely due to the enrichment of resistance associated residues observed in loop D. Predictive analysis indicated that combination of BG18, N6 and PGDM1400 can provide over 95% neutralization coverage at 1μg/mL of contemporary India clade C, an observation found to be distinct to that reported for the Africa clade C viruses. Taken together, we found distinct neutralization patterns and *env* signatures associated with resistance to key bnAbs. Our study highlights that towards achieving clinical effectiveness, both the complementarity of bnAb classes and the regionally relevant HIV forms need to be considered.

**Author summary:** While the development of vaccines to prevent HIV infection remains a global priority, their potential effectiveness is limited by the extraordinarily diversified circulating forms of HIV-1. The prospect of best-in-class bnAbs as potential prevention option has been demonstrated in several studies including the Phase II Antibody Mediated Prevention (AMP) trial; however, to be broadly applicable, bnAbs will need to overcome the substantial variability of HIV *env*. The present study highlights that the contemporary HIV-1 clade C viruses are evolving to be less sensitive to the best-in-class bnAbs and HIV-1 clade C that predominates in India and South Africa vary in their degree of susceptibility to best-in-class clinically relevant bnAbs. This indicates differences in the antigenic properties between globally circulating HIV-1 clade C at a population level. Overall, the outcome of this study highlights the need for periodic assessment of sequence and neutralization profiles of the circulating regionally relevant HIV-1 forms towards prioritizing the bnAb combination suitable for effective intervention.

## Introduction

HIV with complex and evolving diversity [1] remains a global health priority with over 39 million people currently infected globally, 2.4 million of which reside in India making India the third largest HIV epidemic globally [2, 3]. The high genetic variability of globally circulating HIV, both between and within an individual has been a major roadblock in designing an effective preventive intervention despite significant efforts [4]. While antiretroviral therapy has been successful in treatment of HIV and slowing down the spread of the initial pandemic, rising global resistance to available antiretrovirals necessitates expansion of available therapeutics and has reinvigorated efforts to design effective vaccines [5]. In the absence of an efficacious vaccine against HIV, passively administered broadly neutralizing monoclonal antibodies (bnAbs) along with effective antiretroviral drugs (ARV) could play a significant role in reducing incidence in high risk groups and key populations [6]. While efforts towards developing vaccine immunogens capable of inducing broadly neutralizing antibodies (bnAb) are ongoing, the recently conducted Phase 2B Antibody-Mediated Prevention (AMP) efficacy trial (HVTN 704/HPTN 085) demonstrated that a passively administered bnAb could prevent infection by bnAb sensitive viruses [7, 8]. The study also highlighted that combination of bnAbs would be required for optimal coverage of globally circulating HIV-1 subtypes towards capturing those that are resistant to one bnAb class but sensitive to another. HIV-1 clade C, which is the major globally circulating form, also forms the bulk of infections in South Africa and India, although genetic and functional data pertaining to contemporary HIV-1 forms from India are very limited. HIV-1 evolution over time is believed to contribute to changes in *env* sequence, the sole target of bnAbs, that will likely impact the consistencies with the breadth and potency of bnAbs with clinical relevance to effectively tackle currently circulating forms globally [9–11]. This is particularly important as HIV-1 evolution over time is expected to bring in changes in *env* sequence, the sole target of neutralizing antibodies which would impact the degree of susceptibility of contemporary viruses to bnAbs that are relevant for clinical use [9–11].

Previous studies provided evidence of HIV-1 clade B and non-India C viruses becoming increasingly resistant to select bnAbs over time [9, 11–15]. Our previous study using limited historical (obtained prior to 2014) HIV-1 clade C of Indian origin [10] indicated significant variation in their susceptibility to bnAbs and also indicated that evolving viruses were becoming increasingly resistant to key bnAbs such as CAP256-VRC26.25. In the present study, we examined neutralization profiles of the HIV-1 clade C as pseudoviruses encoding full length *env* (*gp160)* of India isolated between 2020-2023 (contemporary) from nine geographically distinct regions in India, compared with that from South Africa and examined the best-in-class bnAbs that would provide optimal neutralization coverage of contemporary India clade C viruses. The *env* sequence diversity and neutralization profiles of contemporary India clade C were also compared with that of South African origin.

## Results

### Phylogenetic profiles of contemporary HIV-1 clade C from different geographical regions of India

We first examined the phylogenetic relationship of the *env* gene of the contemporary viruses from different geographical regions. We obtained unique full length *env* (*gp160)* sequences from 232 individuals from nine geographically distinct sites in India between 2020 and 2023 (**Fig 1A**). These include sequences obtained from ART naïve early seroconverter and from individuals on ART (**Table S1**). The PCR amplified products were processed for high throughput deep sequencing using Oxford Nanopore technology (ONT) and Illumina based NGS platforms to obtain long and short read sequences. A consensus *env* sequence representative of the major circulating variant for each individual was constructed based on multiple alignment of both short and long read sequences to ensure inclusion of the accurate dominant *env* sequences for further analysis. Phylogenetic analysis was performed using these contemporary Indian *env* sequences along with 17 HIV-1 group M reference sequences (hiv.lanl.gov), shown in **Fig.1B**. No region-specific phylogenetic clustering was observed. Interestingly, we identified five subtype A1, one subtype B and four A1/C recombinants **(Table S1)**. Furthermore, *pol* gene sequencing of HIV+ RNA obtained from the therapy naïve individuals showed that 11% of them contain major (>50% variant frequency in deep sequencing data) drug resistance associated mutations (DRM) with reverse transcriptase (RT) associated DRM found to be more prevalent compared to protease (PR) and integrase inhibitor associated DRMs (**Table S1**). Our observation provides evidence of establishment of infection by the drug-resistant HIV-1.

**Fig. 1.**
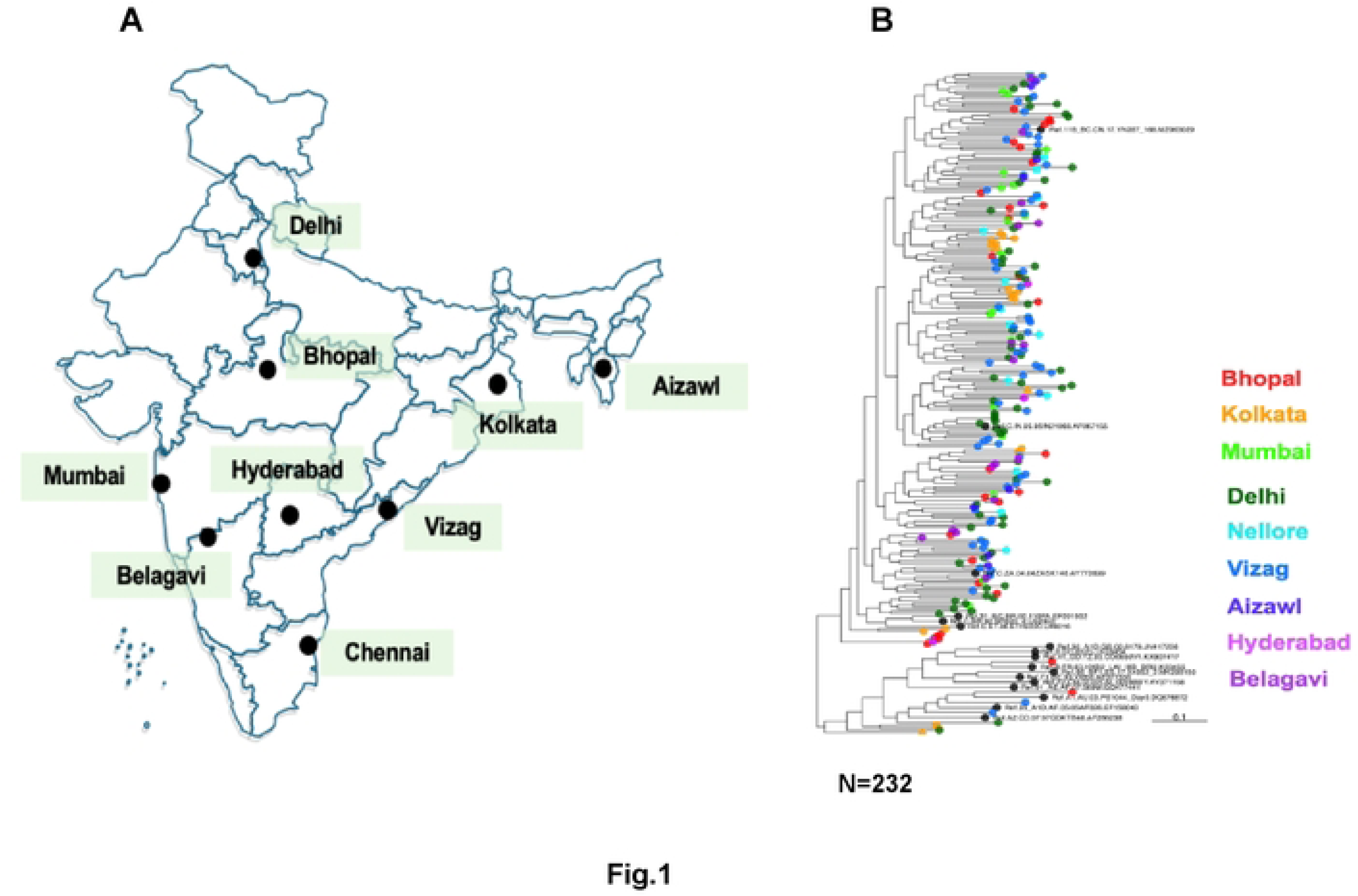
**Phylogenetic relatedness of the contemporary HIV-1 India clade C at the population level**. **A.** Surveillance sites built and samples collected across different geographical sites in India between 2020-2023 **B.** Phylogenetic relatedness of HIV-1 clade C Env proteins representing circulating forms in different geographic regions. Phylogenetic trees were generated for 249 HIV-1 envelope amino acid sequences that included 232 contemporary (obtained between 2020 and 2023) from India and 17 HIV-1 group M reference sequences. These sequences were aligned using MAFFT and the alignment was manually curated in BioEdit v7.2.5. The phylogenetic tree was constructed with IQ-TREE under HIVb model with estimated Ƴ parameters and number of invariable sites. The robustness of the tree topology was further assessed by SH-aLRT as well as 1000 ultrafast bootstrap replicates implemented in IQ-TREE.

### Contemporary HIV-1 India clade C demonstrated significant resistance to V1/V2 directed antibodies compared to those that target CD4 binding and V3 supersites

Next, we examined the neutralization profiles of the contemporary viruses against a large panel of bnAbs with distinct specificities. We randomly selected 115 unique sequences representative all the nine geographically distinct sites in India and prepared pseudoviruses expressing these unique *env* sequences **(Table 1)**, which were further examined against 14 bnAbs of distinct epitope specificities in viral Env. All of these envelopes were found to be CCR5 tropic (**Table 1**). As shown in **Fig. 2**, pseudoviruses expressing contemporary clade C *envs* showed broad sensitivity to bnAbs targeting CD4bs and V3 glycan supersite, compared to ones that target V1/V2 apex. Among bnAbs targeting CD4bs directed bnAbs, N6, 1-18 and VRC07 showed >90% of viruses neutralized with IC50/IC80 values <25μg/mL respectively and over 78% and 65% of the panel viruses neutralized with IC50/IC80 values <1μg/mL respectively) **(Fig.2; Table S2)**. Among the V3 glycan supersite directed bnAbs, 10-1074 and BG18 were observed to demonstrate maximal neutralization breadth (>80% and >70% of the panel viruses neutralized with IC50 and IC80 values <25μg/mL by these bnAbs while over 81% and 74% of the panel viruses neutralized with IC50 and IC80 values <1μg/mL). Amongst all the bnAbs examined, N6 (demonstrated >94% coverage with IC80 of 0.44 μg/mL) and 10-1074 (demonstrated >80% coverage with IC80 of 0.63 μg/mL) were found to be most broad while BG18 (IC80 of 0.29 μg/mL) was found to be most potent. While we identified several clade C viruses with class-specific resistance (**Fig.S1**), V1/V2 directed bnAb class-specific resistant viruses were relatively common **(Table S2)**. A total of 45% and 40% contemporary viruses were found to be resistant to CAP256-VRC26.25 and PGDM1400 respectively. We also identified viruses with resistance to best-in-class bnAbs that target the CD4bs (VRC01, 3BNC117, VRC07, N6 and 1-18) and V3 supersite (PGT121, BG18, 10-1074). Specifically, we identified *env* sequence isolated from four unique donors, which when expressed as pseudoviruses (**Table S2; Fig.S1**) demonstrated complete resistance to all the CD4s-directed bnAbs tested in our study; VRC01, VRC07, N6 and 1-18. For V3 glycan supersite directed bnAbs, we identified eight unique donors, *envs* which demonstrated complete resistance to PGT121, 10-1074 and BG18 (**Table S2; Fig.S1**) Moreover, amongst these individuals, we identified *env* sequences from six individuals which when expressed as pseudovirus demonstrated very broad resistance to the majority of the class-specific bnAbs tested in this study (**Table 2**).

**Fig. 2.**
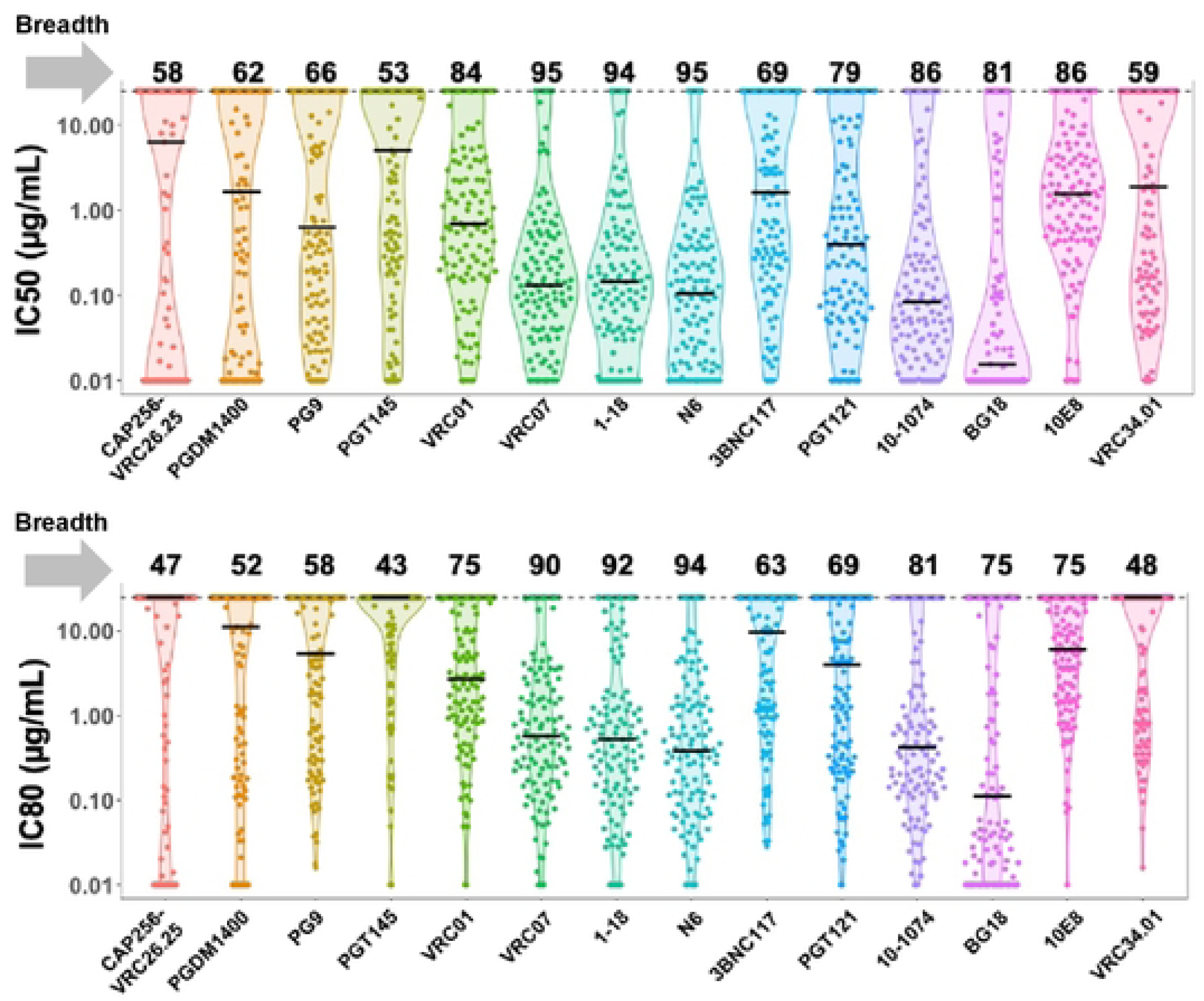
**Neutralization profiles of contemporary HIV-1 India clade C to best-in-class existing bnAbs**. Pseudoviruses expressing 115 contemporary *envs* obtained from individuals representing nine geographically distant regions in India and comprising distinct risk groups were assessed for their degree of susceptibility to 14 bnAbs as indicated having distinct epitope specificities on viral Env. IC50 and IC80 refers to the IgG concentrations [μg/mL] at which pseudoviruses demonstrated 50% and 80% neutralizations respectively. Pseudoviruses that were not neutralized up to 25μg/mL of IgG were considered as resistant viruses. Neutralization assay was carried out at least 3 times in duplicates and average was used to plot the graph. Neutralization breadth of each bnAb expressed as percent neutralization by IgG up to 25μg/mL are shown on top of each graph (upper and lower panel).

**Table 1.** Source, coreceptor usage and other properties of contemporary HIV-1 clade C functional clones.

|  | *env (gp160)* | Subtype | Region | Source | Vector | Coreceptor usage |
| --- | --- | --- | --- | --- | --- | --- |
| 1 | TSG-EHI6 | A1 | Nellore | PLASMA | pCDNA3.1 | CCR5 |
| 2 | TSG-EHI9 | C | Nellore | PLASMA | pCDNA3.1 | CCR5 |
| 3 | TSG-EHI28 | C | Nellore | PLASMA | pCDNA3.1 | CCR5 |
| 4 | TSG-EHI32 | C | Nellore | PLASMA | pCDNA3.1 | CCR5 |
| 5 | TSG-EHI38 | C | Nellore | PLASMA | pCDNA3.1 | CCR5 |
| 6 | TSG-EHI39 | C | Nellore | PLASMA | pCDNA3.1 | CCR5 |
| 7 | TSG-EHI41 | C | Nellore | PLASMA | pCDNA3.1 | CCR5 |
| 8 | TSG-EHI42 | C | Nellore | PLASMA | pCDNA3.1 | CCR5 |
| 9 | TSG-EHI44 | C | Nellore | PLASMA | pCDNA3.1 | CCR5 |
| 10 | TSG-EHI45 | C | Nellore | PLASMA | pCDNA3.1 | CCR5 |
| 11 | TSG-EHIPre8 | C | Nellore | PLASMA | pCDNA3.1 | CCR5 |
| 12 | TSG-EHI13D6 | C | Nellore | PLASMA | pCDNA3.1 | CCR5 |
| 13 | TSG-EHI17B14 | C | Nellore | PLASMA | pCDNA3.1 | CCR5 |
| 14 | TSG-EHI8 | C | Nellore | PLASMA | pCDNA3.1 | CCR5 |
| 15 | TSG-EHI14 | C | Nellore | PLASMA | pCDNA3.1 | CCR5 |
| 16 | TSG-EHI40-C18 | C | Nellore | PLASMA | pCDNA3.1 | CCR5 |
| 17 | TSG-EHI11 | C | Nellore | PLASMA | pCDNA3.1 | CCR5 |
| 18 | TSG-EHI21 | C | Nellore | PLASMA | pCDNA3.1 | CCR5 |
| 19 | TSG-EHI26 | A1 | Nellore | PLASMA | pCDNA3.1 | CCR5 |
| 20 | TSG-EHI30 | C | Nellore | PLASMA | pCDNA3.1 | CCR5 |
| 21 | TSG-EHI18 | C | Nellore | PLASMA | pCDNA3.1 | CCR5 |
| 22 | TSG-EHI22 | C | Nellore | PLASMA | pCDNA3.1 | CCR5 |
| 23 | TSG-EHI29 | C | Nellore | PLASMA | pCDNA3.1 | CCR5 |
| 24 | TSG-EHI35 | C | Nellore | PLASMA | pCDNA3.1 | CCR5 |
| 25 | TSG-EHI37 | C | Nellore | PLASMA | pCDNA3.1 | CCR5 |
| 26 | TSG-EHI7 | C | Nellore | PLASMA | pCDNA3.1 | CCR5 |
| 27 | TSG-EHI12 | C | Nellore | PLASMA | pCDNA3.1 | CCR5 |
| 28 | TSG-EHI20 | C | Nellore | PLASMA | pCDNA3.1 | CCR5 |
| 29 | TSG-EHI23 | C | Nellore | PLASMA | pCDNA3.1 | CCR5 |
| 30 | TSG-EHI27 | C | Nellore | PLASMA | pCDNA3.1 | CCR5 |
| 31 | TSG-EHI33 | C | Nellore | PLASMA | pCDNA3.1 | CCR5 |
| 32 | TSG-EHI55 | C | Nellore | PLASMA | pCDNA3.1 | CCR5 |
| 33 | TSG-EHI60 | C | Nellore | PLASMA | pCDNA3.1 | CCR5 |
| 34 | TSG-EHI61 | C | Nellore | PLASMA | pCDNA3.1 | CCR5 |
| 35 | TSG-EHI62 | C | Nellore | PLASMA | pCDNA3.1 | CCR5 |
| 36 | TSG-EHI63 | C | Nellore | PLASMA | pCDNA3.1 | CCR5 |
| 37 | TSG-EHI25 | C | Nellore | PLASMA | pCDNA3.1 | CCR5 |
| 38 | TSG-EHI50 | C | Nellore | PLASMA | pCDNA3.1 | CCR5 |
| 39 | TSG-EHIPre15 | C | Nellore | PLASMA | pCDNA3.1 | CCR5 |
| 40 | TSG-EHI51 | C | Nellore | PLASMA | pCDNA3.1 | CCR5 |
| 41 | TSG-EHI16 | C | Nellore | PLASMA | pCDNA3.1 | CCR5 |
| 42 | TSG-EHI36 | C | Nellore | PLASMA | pCDNA3.1 | CCR5 |
| 43 | TSG-EHI34 | C | Nellore | PLASMA | pCDNA3.1 | CCR5 |
| 44 | TSG-EHI53 | C | Nellore | PLASMA | pCDNA3.1 | CCR5 |
| 45 | TSG-EHI57 | C | Nellore | PLASMA | pCDNA3.1 | CCR5 |
| 46 | TSG-EHI58 | C | Nellore | PLASMA | pCDNA3.1 | CCR5 |
| 47 | TSG-EHI59 | C | Nellore | PLASMA | pCDNA3.1 | CCR5 |
| 48 | TSG21Y02A0012 | C | Delhi | PLASMA | pCDNA3.1 | CCR5 |
| 49 | TSG21Y02E0018-D18 | C | Delhi | PLASMA | pCDNA3.1 | CCR5 |
| 50 | TSG21Y02E0024-D24 | C | Delhi | PLASMA | pCDNA3.1 | CCR5 |
| 51 | TSG22Y02E0035-DE35 | C | Delhi | PLASMA | pCDNA3.1 | CCR5 |
| 52 | TSG22Y02E0037-DE37 | C | Delhi | PBMC | pCDNA3.1 | CCR5 |
| 53 | TSG21Y02E0011-D11 | C | Delhi | PLASMA | pCDNA3.1 | CCR5 |
| 54 | TSG21Y02E0014-D14 | C | Delhi | PLASMA | pCDNA3.1 | CCR5 |
| 55 | TSG21S01A0008_STMART07 | C | Kolkata | PLASMA | pCDNA3.1 | CCR5 |
| 56 | TSG21S01A0013-STMART12 | C | Kolkata | PLASMA | pCDNA3.1 | CCR5 |
| 57 | TSG21S01A0003-STMART02 | C | Kolkata | PLASMA | pCDNA3.1 | CCR5 |
| 58 | TSG21S01A0014-STM13ART | C | Kolkata | PLASMA | pCDNA3.1 | CCR5 |
| 59 | TSG21S01E0017-STM16E | C | Kolkata | PLASMA | pCDNA3.1 | CCR5 |
| 60 | TSG21S01E0001-STM01E | C | Kolkata | PLASMA | pCDNA3.1 | CCR5 |
| 61 | TSG21S01A0002-STMART01 | C | Kolkata | PBMC | pCDNA3.1 | CCR5 |
| 62 | TSG21S01E0033-C33 | C | Kolkata | PBMC | pCDNA3.1 | CCR5 |
| 63 | TSG21S01E0018-STM17E | C | Kolkata | PLASMA | pCDNA3.1 | CCR5 |
| 64 | TSG21S01A0004-C4 | C | Kolkata | PLASMA | pCDNA3.1 | CCR5 |
| 65 | TSG21S01A0005-C5 | C | Kolkata | PLASMA | pCDNA3.1 | CCR5 |
| 66 | TSG21N01N017 | C | Mumbai | PLASMA | pCDNA3.1 | CCR5 |
| 67 | TSG21N01N011-C10 | C | Mumbai | PLASMA | pCDNA3.1 | CCR5 |
| 68 | TSG21N01N029 | C | Mumbai | PLASMA | pCDNA3.1 | CCR5 |
| 69 | TSG21N01N018 | C | Mumbai | PLASMA | pCDNA3.1 | CCR5 |
| 70 | TSG21N01S028 | C | Mumbai | PLASMA | pCDNA3.1 | CCR5 |
| 71 | TSG21N01N010 | C | Mumbai | PLASMA | pCDNA3.1 | CCR5 |
| 72 | TSG21N01N014 | C | Mumbai | PLASMA | pCDNA3.1 | CCR5 |
| 73 | TSG21N01N023 | C | Mumbai | PLASMA | pCDNA3.1 | CCR5 |
| 74 | TSG21N01N031 | C | Mumbai | PLASMA | pCDNA3.1 | CCR5 |
| 75 | TSG21N01F003 | C | Mumbai | PLASMA | pCDNA3.1 | CCR5 |
| 76 | TSG21N01N001CM | C | Mumbai | PBMC | pCDNA3.1 | CCR5 |
| 77 | TSG21N01N013 | C | Mumbai | PLASMA | pCDNA3.1 | CCR5 |
| 78 | TSG21N01N015 | C | Mumbai | PLASMA | pCDNA3.1 | CCR5 |
| 79 | TSG21N01N025 | A1C | Mumbai | PLASMA | pCDNA3.1 | CCR5 |
| 80 | TSG21N01N026 | C | Mumbai | PLASMA | pCDNA3.1 | CCR5 |
| 81 | TSG21N01S007 | C | Mumbai | PLASMA | pCDNA3.1 | CCR5 |
| 82 | TSG21N01S027 | C | Mumbai | PLASMA | pCDNA3.1 | CCR5 |
| 83 | TSG21N01S030 | C | Mumbai | PLASMA | pCDNA3.1 | CCR5 |
| 84 | TSG21N01S100 | C | Mumbai | PLASMA | pCDNA3.1 | CCR5 |
| 85 | TSG21N01S055 | C | Mumbai | PLASMA | pCDNA3.1 | CCR5 |
| 86 | TSG21N01S062 | C | Mumbai | PLASMA | pCDNA3.1 | CCR5 |
| 87 | TSG22Y03E0010-B10 | C | Bhopal | PLASMA | pCDNA3.1 | CCR5 |
| 88 | TSG22Y03A0015-B15 | A1 | Bhopal | PBMC | pCDNA3.1 | CCR5 |
| 89 | TSG22Y03A0022-B22 | C | Bhopal | PLASMA | pCDNA3.1 | CCR5 |
| 90 | TSG22Y03A0031-B31 | C | Bhopal | PLASMA | pCDNA3.1 | CCR5 |
| 91 | TSG22Y03A0032-B32 | C | Bhopal | PLASMA | pCDNA3.1 | CCR5 |
| 92 | TSG22Y03E0019-B19 | C | Bhopal | PLASMA | pCDNA3.1 | CCR5 |
| 93 | TSG22Y03E0023-B23 | C | Bhopal | PBMC | pCDNA3.1 | CCR5 |
| 94 | TSG23Y07A0012-A12 | C | Aizawl | PBMC | pCDNA3.1 | CCR5 |
| 95 | TSG23Y07A0013-A13 | C | Aizawl | PBMC | pCDNA3.1 | CCR5 |
| 96 | TSG23Y07A0015-A15 | C | Aizawl | PLASMA | pCDNA3.1 | CCR5 |
| 97 | TSG23Y06A012-V12 | C | Vizag | PBMC | pCDNA3.1 | CCR5 |
| 98 | TSG23Y06A015-V15 | C | Vizag | PBMC | pCDNA3.1 | CCR5 |
| 99 | TSG23Y06E001-V1 | C | Vizag | PLASMA | pCDNA3.1 | CCR5 |
| 100 | TSG23Y06A003-V3 | C | Vizag | PBMC | pCDNA3.1 | CCR5 |
| 101 | TSG23Y06A004-V4 | C | Vizag | PBMC | pCDNA3.1 | CCR5 |
| 102 | TSG22Y04A0011-H11 | C | Hyderabad | PBMC | pCDNA3.1 | CCR5 |
| 103 | TSG22Y04A0015-H15 | C | Hyderabad | PBMC | pCDNA3.1 | CCR5 |
| 104 | TSG22Y04A0016-H16 | C | Hyderabad | PBMC | pCDNA3.1 | CCR5 |
| 105 | TSG22Y04A0020-H20 | C | Hyderabad | PBMC | pCDNA3.1 | CCR5 |
| 106 | TSG22Y04A0002-H2 | C | Hyderabad | PBMC | pCDNA3.1 | CCR5 |
| 107 | TSG22Y04A0003-H3 | C | Hyderabad | PBMC | pCDNA3.1 | CCR5 |
| 108 | TSG22Y04A0004-H4 | C | Hyderabad | PBMC | pCDNA3.1 | CCR5 |
| 109 | TSG22Y04A0005-H5 | C | Hyderabad | PBMC | pCDNA3.1 | CCR5 |
| 110 | TSG22Y05A0018-BL18 | A1 | Belagavi | PBMC | pCDNA3.1 | CCR5 |
| 111 | TSG22Y05A0021-BL21 | C | Belagavi | PLASMA | pCDNA3.1 | CCR5 |
| 112 | TSG22Y05A0002-BL2 | C | Belagavi | PBMC | pCDNA3.1 | CCR5 |
| 113 | TSG22Y05E0033-BL33 | C | Belagavi | PBMC | pCDNA3.1 | CCR5 |
| 114 | TSG22Y05E0043-BL43 | C | Belagavi | PBMC | pCDNA3.1 | CCR5 |
| 115 | TSG22Y05A0006-BL6 | C | Belagavi | PBMC | pCDNA3.1 | CCR5 |

**Table 2.**
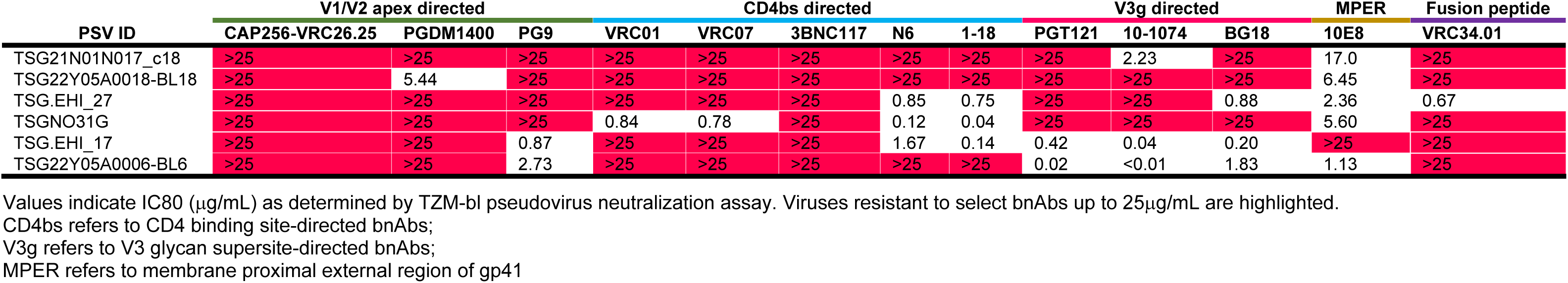
Neutralization profiles of pseudoviruses bearing contemporary *envs* broadly resistant to bnAbs with distinct specificities.

| PSV ID | V1/V2 apex directed |  |  | CD4bs directed |  |  |  |  | V3g directed |  |  | MPER | Fusion peptide |
| --- | --- | --- | --- | --- | --- | --- | --- | --- | --- | --- | --- | --- | --- |
|  | CAP256-VRC26.25 | PGDM1400 | PG9 | VRC01 | VRC07 | 3BNC117 | N6 | 1-18 | PGT121 | 10-1074 | BG18 | 10E8 | VRC34.01 |
| TSG21N01N017_c18 | >25 | >25 | >25 | >25 | >25 | >25 | >25 | >25 | >25 | 2.23 | >25 | 17.0 | >25 |
| TSG22Y05A0018-BL18 | >25 | 5.44 | >25 | >25 | >25 | >25 | >25 | >25 | >25 | >25 | >25 | 6.45 | >25 |
| TSG.EHI_27 | >25 | >25 | >25 | >25 | >25 | >25 | 0.85 | 0.75 | >25 | >25 | 0.88 | 2.36 | 0.67 |
| TSGNO31G | >25 | >25 | >25 | 0.84 | 0.78 | >25 | 0.12 | 0.04 | >25 | >25 | >25 | 5.60 | >25 |
| TSG.EHI_17 | >25 | >25 | 0.87 | >25 | >25 | >25 | 1.67 | 0.14 | 0.42 | 0.04 | 0.20 | >25 | >25 |
| TSG22Y05A0006-BL6 | >25 | >25 | 2.73 | >25 | >25 | >25 | >25 | >25 | 0.02 | <0.01 | 1.83 | 1.13 | >25 |

Values indicate IC80 ( $\mu\text{g/mL}$ ) as determined by TZM-bl pseudovirus neutralization assay. Viruses resistant to select bnAbs up to $25\mu\text{g/mL}$ are highlighted.
CD4bs refers to CD4 binding site-directed bnAbs;
V3g refers to V3 glycan supersite-directed bnAbs;
MPER refers to membrane proximal external region of gp41

We next compared the neutralization profile of the contemporary and historic (isolated prior to 2014) HIV-1 India clade C against CAP256-VRC26.25, PGDM1400, VRC01, VRC07 and PGT121. Contemporary viruses were found to become significantly more resistant to CAP256-VRC26.25 (p<0.005) and more sensitive to PGT121 (p<0.01) when compared with historic viruses (collected before 2014) (**Fig.3A**). While not reaching statistical significance, a trend in contemporary viruses becoming resistant to PGDM1400 and VRC01 was observed. Conversely, a significant increase in sensitivity of contemporary viruses to PGT121 (p<0.0005) was observed. Interestingly, VRC07 was found to demonstrate comparable neutralization of both historical and contemporary Indian clade C, which is in contrast to that observed with contemporary African clade C viruses [9] .

**Fig. 3.**
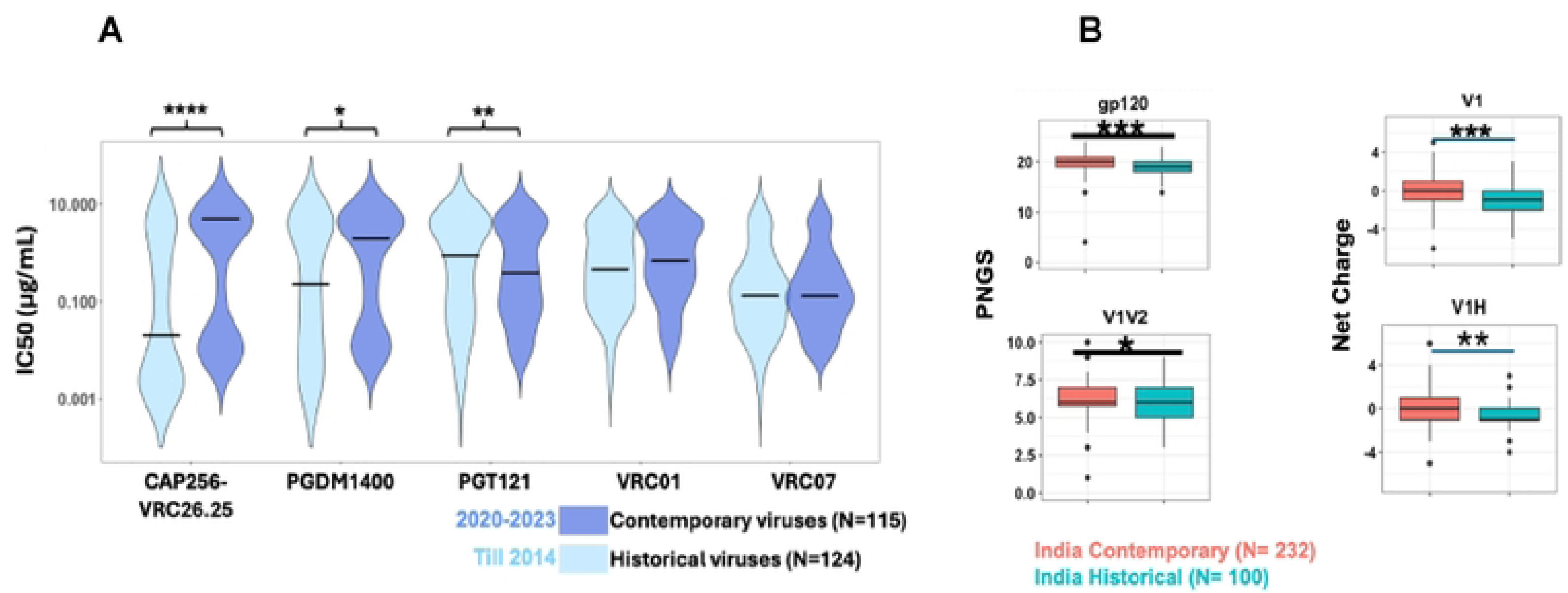
Comparison of neutralization sensitivity to key bnAbs between historic and contemporary India clade C. **A.** Degree of neutralization susceptibility of historical (N=124; obtained before 2014) and contemporary viruses (N=115; obtained between 2020-2023) assessed by pseudovirus neutralization assay. IC50 value of 25 µg/mL was considered as neutralization sensitivity threshold. Statistical analysis to assess significance (P values) of differences in neutralization sensitivity to a given bnAb by pseudoviruses expressing both historical and contemporary *envs* was performed by Mann-Whitney test. Neutralization assay was repeated at least 3 times in duplicates and average was used to plot the graph. **B.** *gp120* variable loop characteristics of historical and contemporary *env* sequences were assessed using the ‘variable characteristics tool’ hosted at the Los Alamos National Laboratory HIV database (LANL-HIVDB, https://www.hiv.lanl.gov/content/sequence/VAR_REG_CHAR/index.html). Potential N linked glycosylation sites prediction was performed with the tool N-Glycosite at LANL-HIVDB (https://www.hiv.lanl.gov/content/sequence/GLYCOSITE/glycosite.html). Statistical significance was assessed by Mann-Whitney test. P values between 0.05-0.01, 0.01-0.001, < 0.001 and <0.0001are depicted as ‘*’, ‘**’, ‘***’ and ‘****’ respectively.

By comparing the *env* sequences of contemporary and historical Indian clade C viruses, we found that overall, they significantly differ in their PNLG site content in *gp120*, specifically in the V1/V2 domain and in their net charge in the V1 hypervariable region. (**Fig.3B**). These features may contribute to reduced sensitivity of contemporary viruses to CAP256-VRC26.25 and PGDM1400, but increased sensitivity to PGT121.

### Neutralization profiles of India and South Africa clade C viruses differ against multiple bnAb classes

We next made a head-to-head comparison of contemporary India and South Africa HIV-1 clade C *envs (gp160)* to examine (a) their phylogenetic relatedness and (b) their sensitivity to bnAbs. For the phylogenetic analysis, we examined 232 and 73 clade C *envs* of Indian and South African origin. Of 73 HIV-1 clade C *env* sequences of African origin, 41 were obtained from individuals enrolled in the FRESH cohort [16] and the rest (32) [9] were obtained from the placebo arm of the phase 2 HVTN 704/HPTN 085 Antibody Mediated Prevention (AMP) prevention trial (South Africa 19, Malawi 7, Zambia 4 and 1 each from Mozambique and Botswana. [7]. As shown in **Fig 4A**, we observed distinct clustering of India and South Africa viruses, consistent with our previous observation [17]. This indicates that *env* genes are genetically distinct and continue to evolve independently in the two geographic regions.

**Fig. 4.**
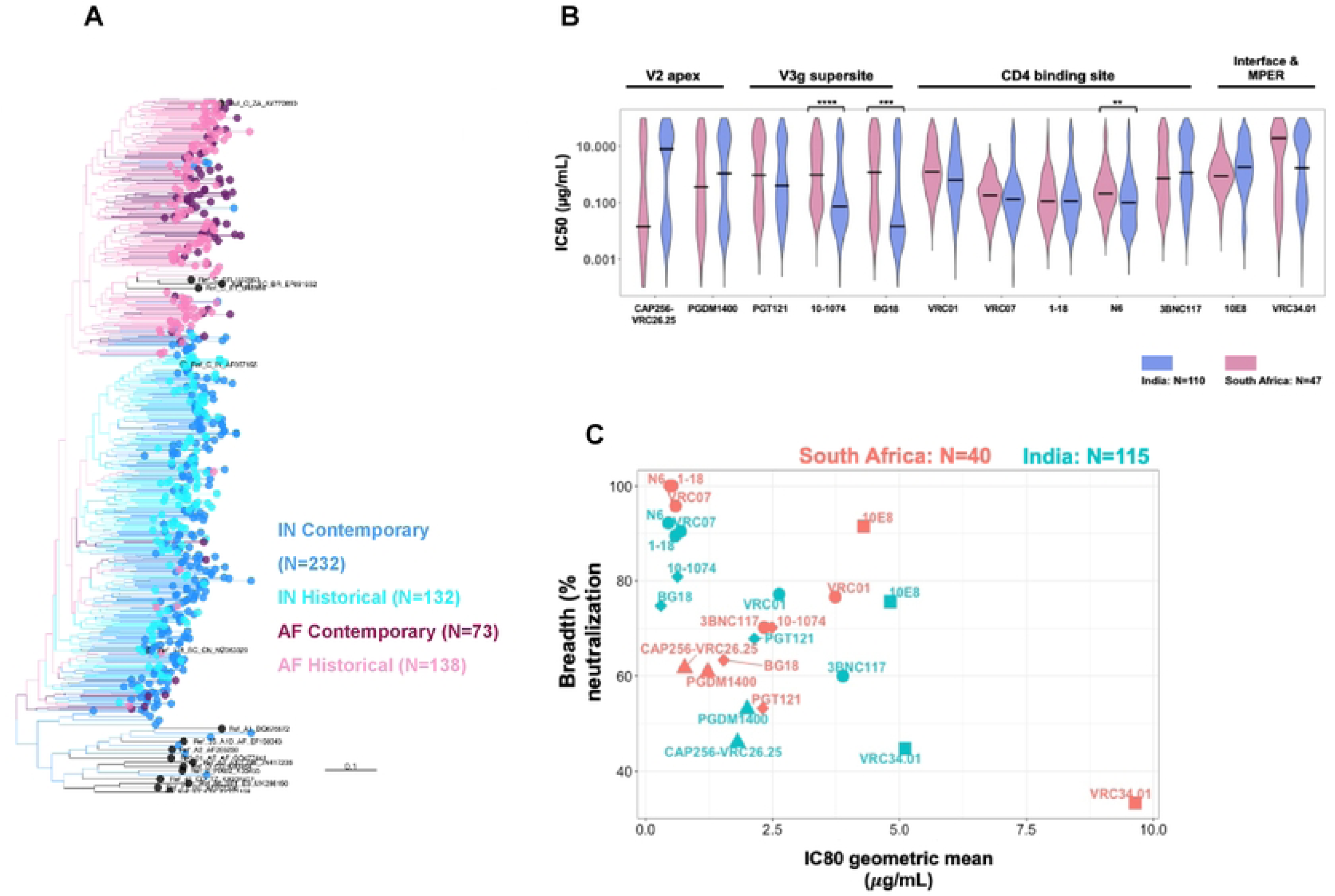
Comparison of phylogenetic and head-to-head neutralization profiles between contemporary India and South African clade C. **A.** Phylogenetic relatedness of *env* genes obtained from contemporary HIV-1 clade C of India (N=232) and Africa (N=73) origins as well as historical India (N=132) and Africa (N=138) origins and 17 HIV-1 group M reference sequences. South Africa clade C *envs* comprised those obtained from FRESH cohort (N=41) and AMP placebo arm (N=32). **B.** Comparison of the degree of neutralization susceptibility of pseudoviruses expressing contemporary HIV-1 clade C *envs* of Indian (N=115) and South African (N=40; AMP placebo arm) origins to 12 best-in-class bnAbs with distinct epitope specificities on viral Env. Env expressed as pseudovirus that showed IC50 value >25 µg/mL against a particular bnAb was considered as resistant. Statistical analysis to assess significance (P values) of differences in neutralization sensitivity to a given bnAb by pseudoviruses expressing *envs* of India and South African origins was assessed by Mann-Whitney test. **C.** Comparison of the magnitude of neutralization sensitivity India and South Africa clade C viruses to select clinically relevant bnAbs. The neutralization breadth of each bnAb tested against India and South Africa clade C envelopes is expressed in the Y-axis as percent neutralization at given concentration of corresponding antibody (IgG) concentration given in X-axis. The values in X-axis are the geometric mean of the IC80 values (μg/mL) calculated for each bnAb. Neutralization assay was carried out in duplicate replicates at least 3 times and average values were used to plot the graph.

We next compared neutralization sensitivity of contemporary Indian (N=115) and South African clade C (N=47, obtained from AMP Placebo group) *envs* against 14 bnAbs described above. We observed significant differences in their neutralization susceptibility to N6 (p<0.05), 10-1074 (p<0.0005) and BG18 (p<0.005) (**Fig.4B),** with Indian HIV-1 clade C being significantly more sensitive to these three bnAbs than African viruses. In general, we observed that except for VRC01 and 3BNC117, comparable neutralization sensitivity to VRC07, 1-18 and N6 observed between Indian and South Africa clade C viruses. However, India clade C viruses demonstrated increased sensitivity to the V3 glycan supersite directed bnAbs examined. Notably, both India and African contemporary clade C viruses were found to show poor susceptibility to CAP256-VRC26.25 and PGDM1400 compared to other bnAbs. However, the degree of resistance to CAP256-VRC26.25 and PGDM1400 were found to differ between India and South Africa clade C (**Fig.4C**.). Furthermore, while both Indian and African clade C viruses are broadly resistant to CAP256-VRC26.25, Indian clade C viruses were found to be more resistant to CAP256-VRC26.25 compared to that observed with African viruses as determined by their mean IC50 values Overall, Indian and African HIV-1 clade C vary significantly in their bnAb neutralization profiles, highlighting the divergence that can occur, even within the same clade.

### Diversity in sequence characteristics that differentiate neutralization sensitive and resistant envelopes

Next, we examined the amino acids in Env that form bnAb contact sites. We created sequence logos to examine the distribution of amino acids and performed statistical tests to assess enrichment of resistance associated signatures. Since the majority of the contemporary viruses were resistant to the V1/V2 directed bnAbs (CAP256-VRC26.25 and PGDM1400), we first examined the distribution of relevant amino acid residues within these epitopes. As shown in **Fig. 5A**, for CAP256-VRC26.25 resistant viruses, we saw signals at positions 160, 166, 169, 170, 200, 332, 632 and 775. We observed significant increases in the frequency of R169, Q170, T332 and decreased frequency of N160, K169 in CAP256-VRC26.25 resistant viruses when compared with the CAP256-VRC26.25 sensitive viruses. For PGDM1400, we observed enrichment of resistance associated residues at positions 160, 169, 172, 275, 332 (**Fig.5B**). Variation in V1/V2 loop length has been shown to modulate sensitivity to V2 apex directed neutralizing antibodies [18–20]. The CAP256-VRC26.25 resistant viruses were also found to have significantly lower net charge in V2 region (P=0.006) compared to their sensitive counterparts (**Fig.5C**). Unlike South African clade C viruses, no significant differences were seen in V1 loop length of CAP256-VRC26.25 sensitive and resistant India clade C viruses (**Fig.5C**; **Fig.S2**), nor was there a significant difference in V2 loop length and net charge between PGDM1400 sensitive and resistant India clade C viruses (**Fig.5C**; **Fig.S2**). For V3 directed bnAbs, we examined residues associated with resistance to PGT121, 10-1074 and BG18. We observed higher variation/entropy within such residues in PGT121 resistant viruses followed by 10-1074 and BG18 resistant viruses **(Fig.S3)**. For PGT121 resistant viruses, significant enrichment and reduction of key residues at position 137, 139, 140, 328, 332, and 334 was observed **(Fig.S3A)**. In 10-1074 resistant viruses, there was significant enrichment of A137, E322, K328, Y330, T332 and N334 compared with sensitive viruses **(Fig.S3A)**. While most contemporary viruses were potently neutralized by BG18, the few that showed resistance were significantly enriched for Y330 and N/T330 **(Fig.S3A)**. For PGT121 resistant viruses, we observed low net charge in the V1/V2 hypervariable region compared to the PGT121 sensitive viruses (P=0.04). We also observed significant differences in V4 loop length (P=0.02) and net charge in the V1 loop (P=0.04) in the 10-1074 resistant viruses (**Fig.S3B**). With respect to BG18 resistant viruses, we observed significant differences in the net charges in the V1 loop (P=0.03), V4 loop length (P=0.01) and PNLG content in V4 loop (P=0.01) when compared with BG18 sensitive viruses (**Fig.S3B**). Amongst CD4bs directed bnAbs examined, contemporary viruses showed least susceptibility to 3BNC117 (26.08% were found to be resistant) followed by VRC01 (23.47%), 1-18 (10.43%), VRC07 9.56%) and N6 (7.82%) (**Fig.8A**). We observed significant enrichment of E279, S280, R282, F318, V371, E455 and significant reduction in the occurrence of N280, Y318, S365, I371, R456, G459, G471 in viruses resistant to 3BNC117 **(Fig.S4)**. Overall, in VRC01 resistant viruses, we observed enrichment of aspartic (D) and glutamic acid (E) residues at position 97 in C1 region of the envelope inner domain, polymorphisms at 279 and 281 positions in the loop D and enrichment of glutamic acid (E) and/or leucine (L) at 455 position, tryptophan (W) at 456 position, aspartic acid (D) at asparagine (N) at the positions 455, 456 and 474 positions in the β23/loop-β24/V5 region of the viral Env protein. Similarly, in 3BNC117 resistant viruses, we observed significant polymorphism at positions 279, 280, 282, 318, 371, 455, 459 and 471 on viral Env protein that are associated with modulation of sensitivity to 3BNC117 (www.hiv.lanl.gov). Interestingly, we observed enrichment of several amino acid residues at positions 279, 280, 281, 355, 365, 456, 459, 463 and 471 in the N6 resistant viruses **(Fig.S4)** around CD4bs region. Significant differences observed between the variable region of viruses sensitive and resistant to VRC01, VRC07 and N6 are shown in **Fig.S5**.

**Fig. 5.**
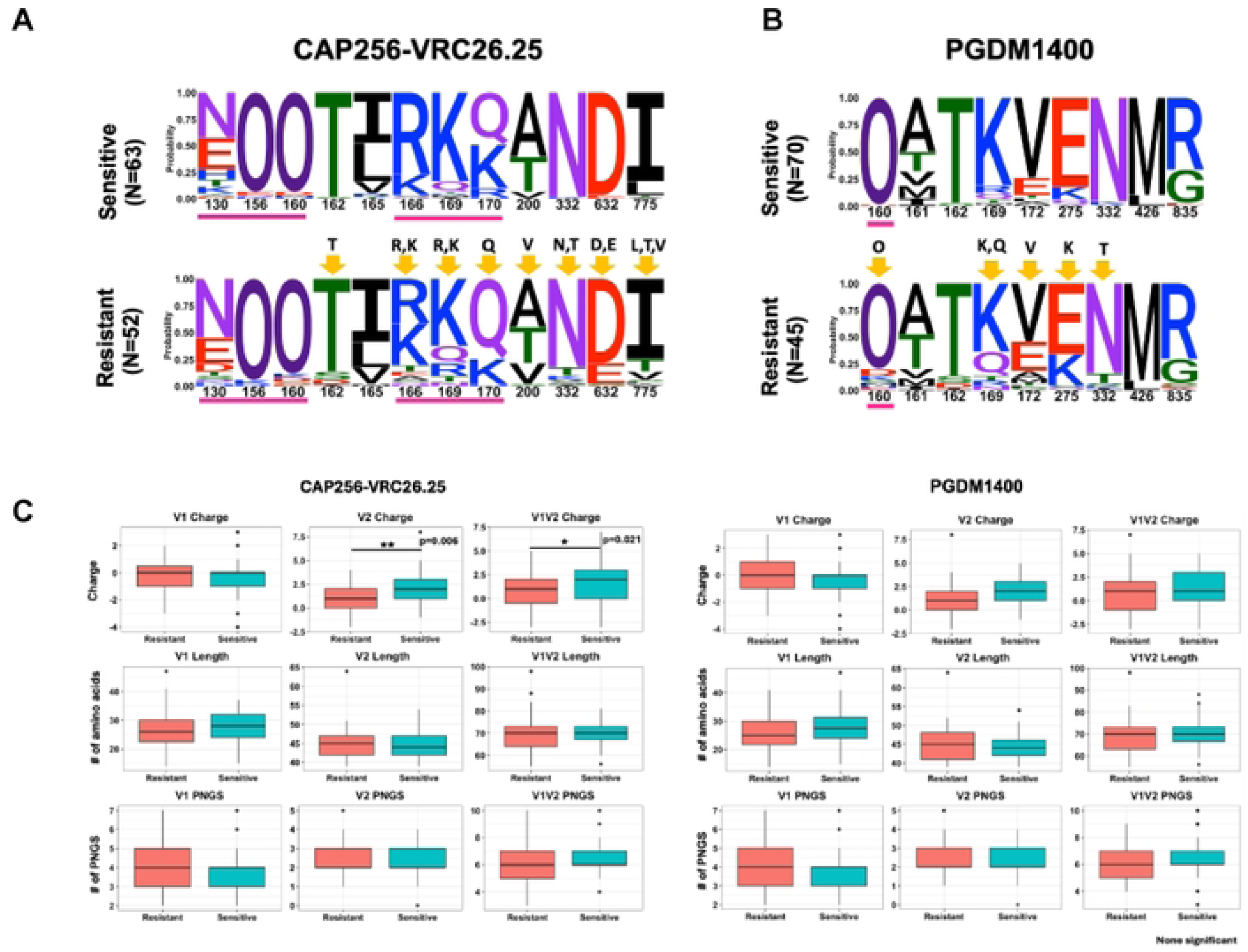
Diversity in *gp120* sequence features and contact sites polymorphism between contemporary India clade C Envs sensitive and resistant to V/1/V2 apex clinically relevant bnAbs. Frequency of contact sites associated with CAP256-VRC26.25 and PGDM1400 sensitivity were compared between CAP256-VRC26.25 sensitive and resistant pseudoviruses **(A)** and PGDM1400 sensitive and resistant viruses **(B)**. The gp160 position (based on HXB2 numbering) of the key amino acids in the sequence logo are shown in X-axis and their relative abundance expressed as probability in Y axis. O has been used to differentiate potential N linked glycosylated Asparagine from potentially unglycosylated Asparagine (N). Residues underscored in purple line are direct Ab contact sites. Residues showing statistically significant changes in abundance following a Fisher’s exact test are highlighted with yellow arrows. **C**. Variable loop length, PNGs and net charges of sensitive and resistant envelopes.

When we compared the sequence features of the contemporary HIV-1 clade C *env* of Indian and South African origins, we observed a significant difference in their *gp160* (both *gp120* loop and *gp41)* lengths **(Fig.6A)**. In particular, we found a significant difference in the V4 loop length between the contemporary viruses from these two geographically distinct regions with longer loops observed with Indian contemporary viruses. Moreover, contemporary India and Africa clade C *envs* also significantly differed in their V1/V2 net charge and PNLGs in the V4 loop (**Fig.6A**). We compared the *env* sequences of India and Africa viruses that showed resistance to V1/V2 directed bnAbs. For CAP256-VRC26.25 and PGDM1400 resistant viruses, we found differences at sites 160, 166, 169 and 170 (**Fig.6B**). Interestingly, except for differences in net charge in V2 loop between PGDM1400 resistant viruses of India and South Africa clade C, no differences in loop lengths and PNGs observed between CAP256-VRC26.25 and PGDM1400 resistant India and South Africa clade C viruses suggestive of similar mechanisms of resistance across the two regions (**Fig.S6**). We also observed differences in frequencies of contact residues targeted by CD4bs (N6) and V3 glycan directed (PGT121, 10-1074, BG18) bnAbs between India and South Africa clade C viruses **(Fig.S7)** which may explain the differences in their sensitivity (**Fig. 4B)**. While the above analysis was carried out using the *env* sequences which were expressed and tested against the selected bnAbs as pseudoviruses, we analyzed additional contemporary clade C *env* sequences (not used for preparing pseudoviruses) isolated from nine geographically distinct regions of India (as described above) and from South Africa (sourced from FRESH cohort). While both datasets displayed enrichment of resistant signatures for CAP256-VRC26.25 bnAb at positions 165, 166 and 169, statistically significantly different enrichment of resistant signature was observed at residue position 166 **(Fig.S8)**. Analysis of PGDM1400 contact residues indicated a trend of differential abundance at position 160 with significant differences at positions 130 and 161 as well as 211. N332, the target site for PGT121, 10-1074 and BG18 bnAbs was more conserved in Indian sequences compared to those from South Africa **(Fig.S8)**. Overall, our data indicated that the differential sensitivity of India and Africa clade C contemporary viruses bnAb classes is associated with distinct sequence features including those in bnAb contact residues.

**Fig. 6.**
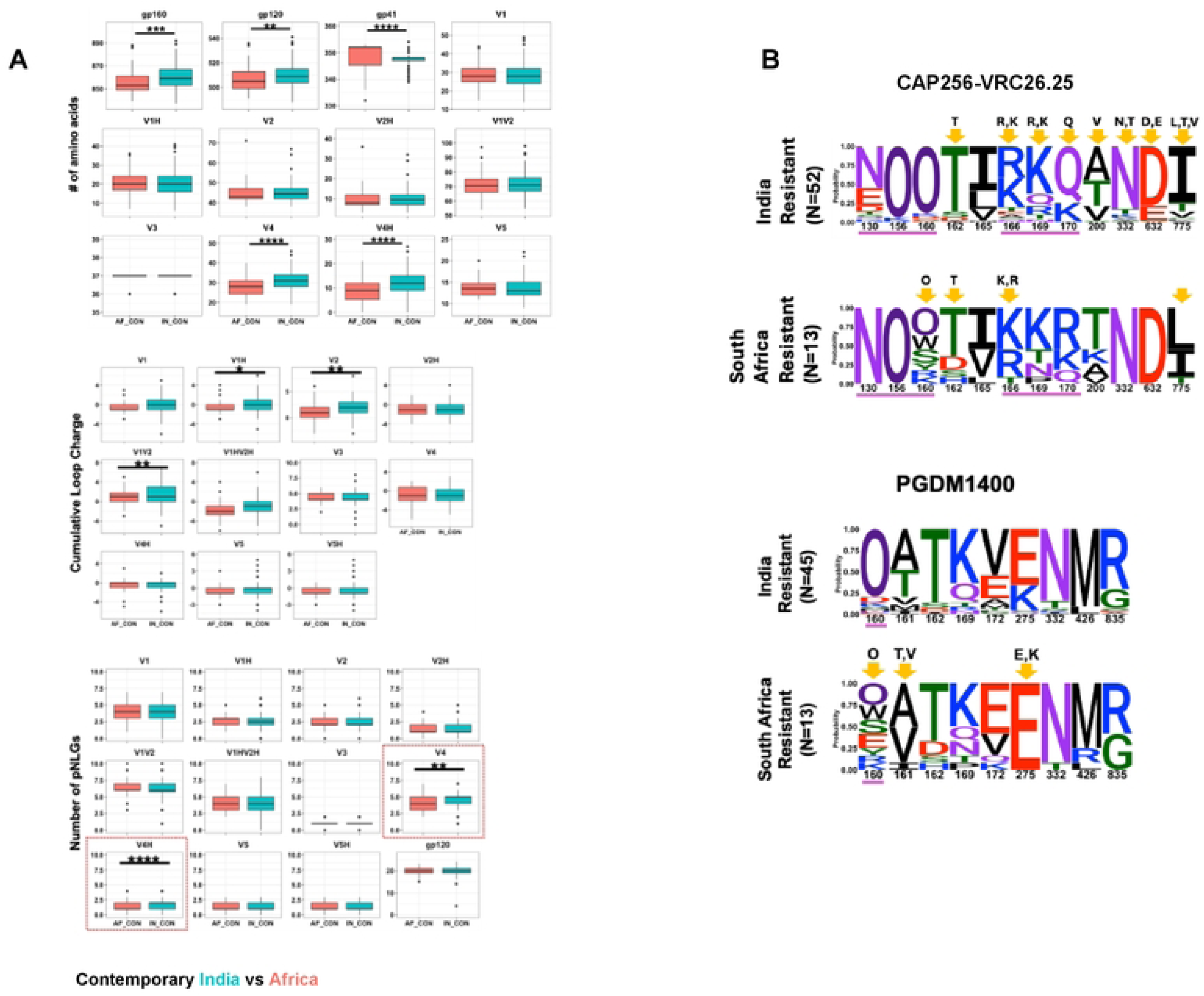
Comparison of *env* sequence features of contemporary India and South Africa clade C viruses. **A.** The amino acid sequences of complete *envs* (*gp120 and gp41*) of India and South Africa contemporary HIV-1 clade C were analyzed to compare their average variable loop lengths, PNLGs and net charges in *gp120* as well as the length of gp41. These are analyzed using ‘Variable region characteristics’ tool available at the Los Alamos HIV database (https://www.hiv.lanl.gov/content/sequence/VAR_REG_CHAR/index.html) and N-Glycosite (https://www.hiv.lanl.gov/content/sequence/GLYCOSITE/glycosite.html). **B.** Comparison of key amino acid residues on India and South Africa clade C *envs* that are linked with CAP256-VRC26.25 and PGDM1400 resistance is shown in Sequence logos. The statistically significant enrichment of key residues for viruses sensitive and resistant to CAP256-VRC26.25 and PGDM1400 are shown in Y-axis. O has been used to differentiate potential N linked glycosylated Asparagine from potentially unglycosylated Asparagine (N). Amino acid residues underscored in purple line are direct Ab contact sites for respective bnAbs. Residues showing statistically significant changes in abundance following a Fisher’s exact test are highlighted with yellow arrows

### Viruses resistant to V1/V2 directed antibodies remain well neutralized by CD4bs directed antibodies

We next examined the ability of other bnAbs to neutralize V1/V2 directed bnAb resistant viruses. As shown in **Fig.7**, CAP256-VRC26.25 resistant viruses were found to be best neutralized by CD4bs directed bnAbs (82.25% breadth) over the V3 directed bnAbs (74.19%). Amongst the CD4bs directed bnAbs, N6 and 1-18 demonstrated best breadth (91.93%) and 3BNC117 was found to be least broad amongst all (58.06%). As for V3 directed bnAbs, CAP256-VRC26.25 resistant viruses were found to be best neutralized by 10-1074 (80.64%), followed by BG18 (75.80%) and PGT121 (66.13%). When we analyzed viruses that demonstrated complete resistance to all the V1/V2 apex directed bnAbs, we again found that compared to V3 directed bnAbs (65.38% breadth), they are best neutralized by CD4bs directed bnAbs (81.53%) with both N6 and 1-18 demonstrating maximum breadth (92.30% breadth in both), however N6 was found to be more potent with IC50 of 0.35μg/mL over 1-18 with IC50 of 1.01 μg/mL. Our data indicates that while contemporary Indian clade C viruses showed poor susceptibility to V1/V2 directed bnAbs, they remain broadly sensitive to lead CD4bs and V3 specific bnAbs.

**Fig. 7.**
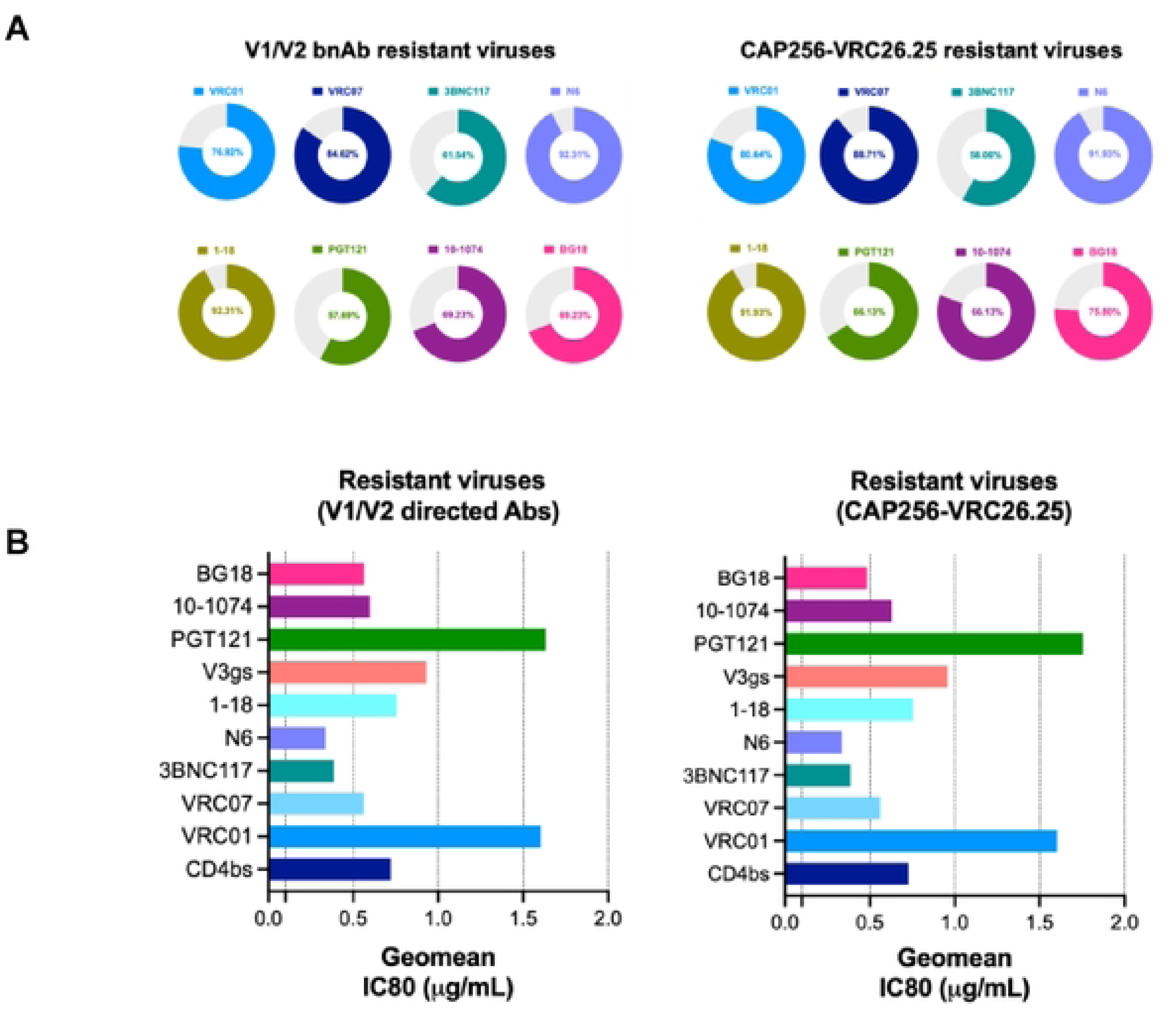
Neutralization of V1/V2 apex bnAb resistant contemporary Indian clade C viruses by CD4bs and V3 glycan supersite directed bnAbs. **A.** Sensitivity of pseudoviruses expressing contemporary Indian clade C *envs* which were fully resistant to all V1/V2 directed bnAbs to CD4bs (VRC01, VRC07, 3BNC117, N6 and 1-18) and V3 glycan supersite (PFT121, 10-1074 and BG18). Left panel shows percent neutralization of pseudoviruses that were resistant to all V1/V2 directed bnAbs tested (CAP256-VRC26.25, PGDM1400, PG9) (N=26) by CD4bs and V3 glycan directed bnAbs. Right panel shows same but only to pseudoviruses resistant CAP256-VRC26.25 resistant envelopes (N=62). Percent neutralization breadth conferred by CD4bs and V3 glycan directed bnAbs was calculated by the number of resistant viruses showed IC80 values <25μg/mL. **B**. Magnitude of neutralization of V1/V2 directed bnAb resistant pseudoviruses conferred by each of the CD4bs and V3 glycan directed bnAbs. The magnitude of virus neutralization equivalent to potency was measured as the lowest geometric mean titer (GMT) of the conferred by each bnAb IgG (μg/mL) that demonstrated 80% neutralization of pseudovirus. Neutralization assay was carried out in duplicate replicates at least 3 times and average values were used to plot the graph.

### VRC01 & 3BNC117 resistant viruses are neutralized by second generation CD4bs bnAbs but with reduced potency

While CD4bs directed antibodies were found to demonstrate best neutralization coverage of the contemporary viruses, we next examined whether resistance to CD4bs bnAbs VRC01 and 3BNC117 conferred decreased sensitivity to second generation CD4bs bnAbs. As shown in **Fig.8A**, amongst all the CD4bs bnAbs tested, contemporary Indian clade C showed greater resistance to VRC01 and 3BNC117 (23.48% and 26.08% respectively) compared to those that showed resistance to VRC07 (9.56%), N6 (7.82%) and 1-18 (10.43%). We next examined the extent of neutralization of VRC01 and 3BNC117 resistant contemporary viruses by other CD4bs bnAb classes. We observed that N6 neutralized (77.77%) most of the VRC01 resistant viruses, while 1-18 could neutralize (75%) most of the 3BNC117 resistant contemporary viruses (**Fig.8B**). Interestingly, when compared with VRC01 and 3BNC117 sensitive viruses, VRC07, N6 and 1-18 were found to neutralize VRC01 and 3BNC117 resistant viruses with reduced potency by over 2-fold (**Fig.8C**). The reduced potencies could likely be due to enrichment of resistance associated amino acid residues observed when we compared the VRC01 and 3BNC117 sensitive and resistant viruses. Overall, we found that in addition to clinically relevant V1/V2 bnAb resistant viruses, the second generation CD4bs bnAbs such as VRC07, N6 and 1-18 are capable of neutralizing viruses that are resistant to first generation CD4bs bnAbs VRC01 and 3BNC117.

**Fig. 8.**
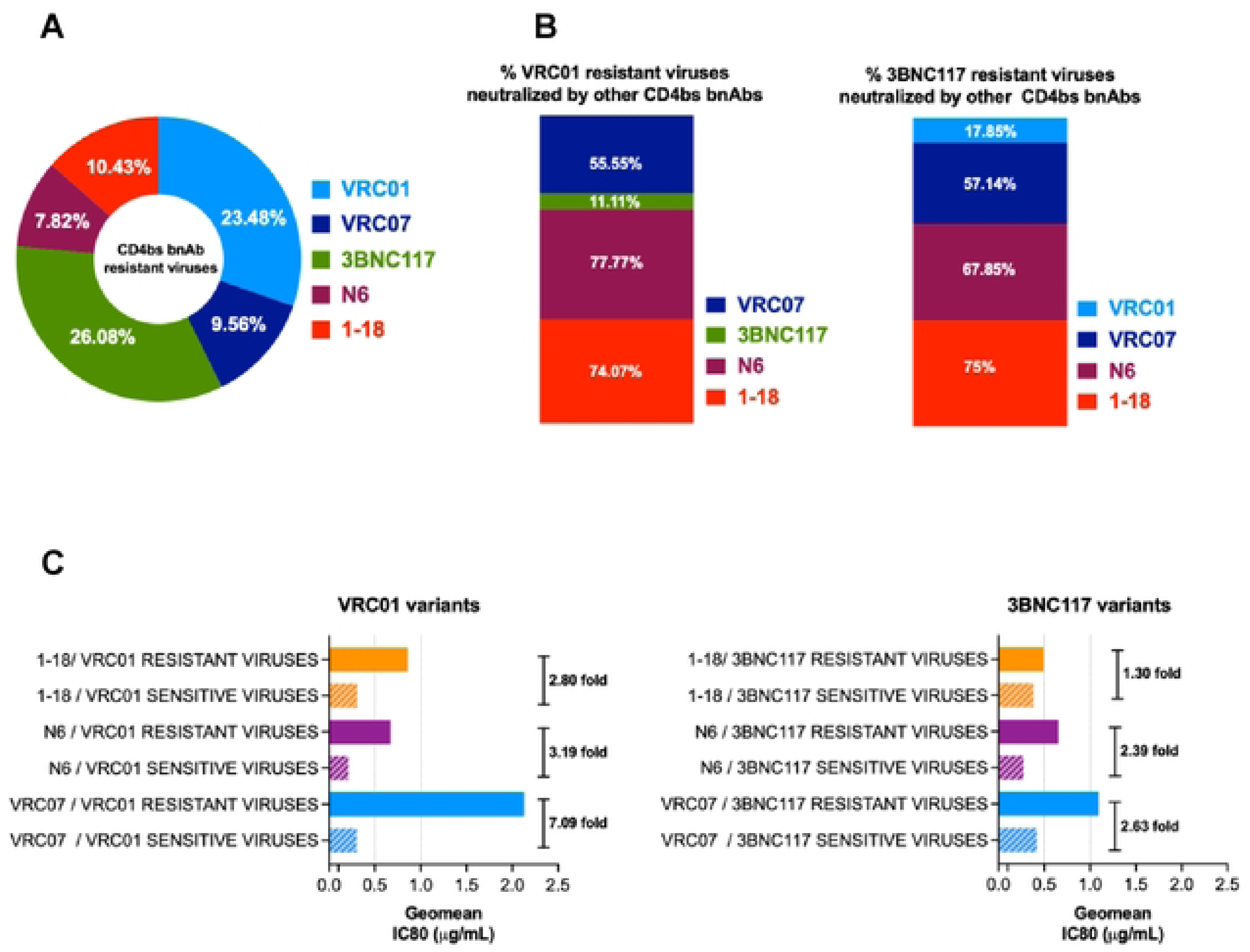
Neutralization efficiency of VRC01 and 3BNC17 resistant contemporary viruses by second generation CD4bs directed bnAbs. **A.** Proportion of contemporary viruses (N=115) that were found to be resistant to first (VRC01 and 3BNC117) and second (VRC07, N6, 1-18) generation CD4bs directed bnAbs. Pseudoviruses with neutralization score of IC80 >25 μg/mL to respective bnAbs were considered resistant. **B.** Proportion of VRC01 and 3BNC117 resistant contemporary pseudoviruses that demonstrated sensitivity to second generation CD4bs bnAbs (VRC07, N6, 1-18). Note that both VRC01 and 3BNC117 resistant viruses were least neutralized by 3BNC117 (11.11%) and VRC01 (17.85%) compared to VRC07, N6 and 1-18, indicating that they viruses resistant to both of them lacks common key residues that are essential for both VRC01 and 3BNC117 for comprehensive neutralization. All the second generation CD4bs bnAbs showed better neutralization (over 50%) with 1-18 demonstrated most (>74%) of VRC01 and 3BNC117 resistant viruses. **C.** Comparison of magnitude of neutralization of VRC01 and 3BNC117 sensitive and resistant viruses by second generation CD4bs bnAbs. Left panel shows the differences in the magnitudes of neutralization of VRC01 sensitive and resistant viruses by all the three CD4bs bnAb (VRC07, N6, 1-18) and right panel same with 3BNC117 sensitive and resistant viruses. The fold difference in magnitude of neutralization was obtained by calculating the average (GMT) of IC80 (μg/mL) for each paired set. GraphPad Prism was used to plot all the graphs.

### Combination of BG18, N6 and PGDM1400 is predicted to provide optimal neutralization of India clade C

Towards identifying the most optimal combination of bnAbs capable of comprehensively neutralizing contemporary clade C viruses, we included bnAbs that demonstrated neutralization breadth >50% with IC80 of <25µg/mL. Also, in order to perform a head-to-head comparison, we assessed the extent of neutralization coverage of the contemporary clade C viruses from Africa (N=40) by the same set of bnAbs. The CombiNAber analysis (https://www.hiv.lanl.gov/content/sequence/COMBINABER/combinaber.html) was carried out for both set of viruses (of India and Africa origins) at the target concentration of 1µg and 10µg/mL respectively using the Bliss-Hill model. At 10 µg/mL, Indian contemporary viruses were observed to be most effectively neutralized by N6 and BG18 (**Fig.9A**). Both of these provided 91% and 72% coverage at the target concentration with potency (IC80) of 0.44 and 0.30 µg/mL respectively. 1-18 and 10-1074 were the next best two CD4bs and V3 directed bnAbs with breadth of 83 and 80 and potency (IC80) of 0.59 and 0.63 µg/mL respectively. With respect to the contemporary clade C viruses from Africa, N6 was observed to be the most effective bnAb with 90% coverage and potency (IC80) of 0.51 µg/mL. PGDM1400 and BG18 were comparably next most effective bnAbs with neutralization breadth of 57.7% and 56.6% and potency (IC80) of 1.59 and 1.54 µg/mL respectively. When we assessed 3 bnAb combination prediction, BG18 + N6 + PGDM1400 appear to provide best neutralization coverage of Indian contemporary viruses with 99.13% breadth with IC80 predicted to be at 0.03 µg/mL. However, the coverage drops to 79% when considering at least two active bnAbs. For the HIV-1 clade C from South Africa, the combination of BG18 + PGDM1400 + 1-18 appear to be the best combination with 100% coverage at 0.03 ug/mL IC80. The coverage drops to mere 81.81% when considering at least two active bnAbs. At 1 µg/mL, Indian contemporary clade C viruses appeared to be most effectively neutralized by BG18 and N6 (**Fig.9B**) as above. However, they showed 64 and 66% neutralization coverage at the target concentration with potency (IC80) of 0.30 and 0.44 µg/mL respectively. Similarly, as with 10µg/mL concentration, 1-18 and 10-1074 were the next best bnAbs found with predicated neutralization coverage of 64% and 66% and potency (IC80) of 0.59 and 0.63 µg/mL respectively. For the contemporary clade C viruses from South Africa, the most effective single mAbs were found to be 1-18 and BG18. These two bnAbs were observed to provide 65 and 43% coverage respectively, with potency of 0.53 and 1.54 ug/mL respectively. When we assessed neutralization coverage by three antibody combination, BG18 + N6 + PGDM1400 observed to provide 93.91% coverage of Indian contemporary clade C viruses at IC80 of 0.03 ug/mL. This neutralization coverage, however drops to 58% when at least two active bnAbs were considered. Conversely, we observed BG18 + PGDM1400 + 1-18 combination to provide 95.45% coverage of African contemporary viruses at IC80 of 0.03 ug/mL and which drops significantly to 45.45% when considering at least two active bnAbs. Overall, our predictive data indicates that no combination could provide 100% coverage for the clade C viruses from India and African origin at any of the considered target concentrations. The data further indicated that while a combination of V3, V2 apex and CD4bs directed bnAbs was effective across both regions, the clade C viruses from India and Africa are distinctly sensitive to different bnAbs of clinical relevance.

**Fig. 9.**
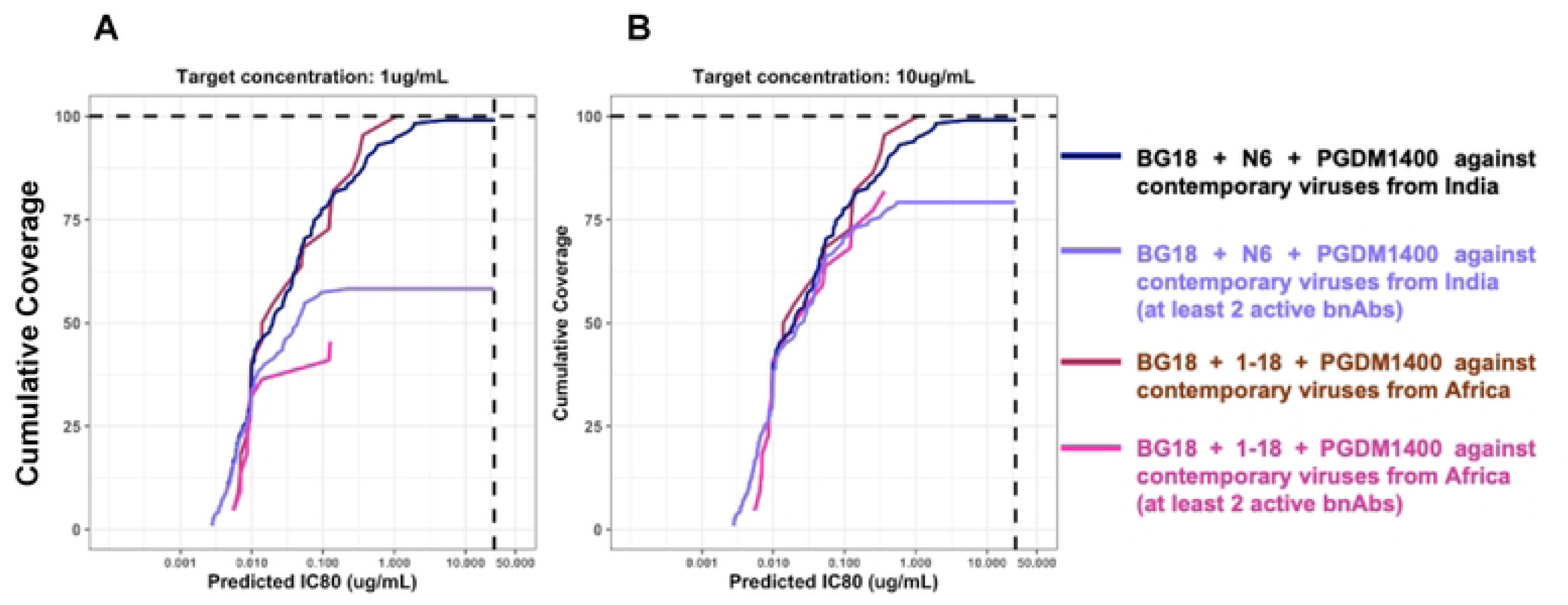
Predictive neutralization coverage of contemporary India clade C viruses by clinically relevant bnAbs. Cumulative neutralization coverage of pseudoviruses carrying contemporary HIV-1 clade C *envs* by bnAb combination was assessed using CombiNAber tool using the Bliss-Hill statistical model. (https://www.hiv.lanl.gov/content/sequence/COMBINABER/combinaber.html). CombiNAber analysis of 115 contemporary viruses from India against BG18 + N6 + PGDM1400 and 45 Contemporary viruses from Africa against BG18 + 1-18 + PGDM1400 as well as the same Combinations with at least 2 active bnAbs have been plotted for target bnAb concentrations of 1ug/mL and 10ug/mL respectively. Predicted IC80 (μg/mL) combinations have been plotted on the X axis while the cumulative breadth of the viruses has been depicted on the Y axis.

## Discussion

While HIV-1 clade C is the major globally circulating form, evolutionary pattern may vary across different geographical regions representing ethnically diversified populations that may contribute to differential susceptibility to class-specific bnAbs. For example, there has been significant association between HIV evolution at population level and increased resistance to serum and bnAb mediated neutralization observed in HIV-1 clade B infected individuals [11–13]. Moreover, intra-clade diversity has been predicted to have better neutralization advantage in geographical regions with lower viral diversity compared to regions with substantial intra-clade diversities[21]. Therefore, it is unclear whether same combination of select bnAbs would stand fit to comprehensively provide neutralization coverage of the globally circulating and evolving HIV at the population level. It is therefore important to understand whether globally evolving HIV at a geographically and ethnically distinct population level can influence antigenic properties. Little information is available for contemporary HIV-1 clade C viruses predominantly circulating across India.

In the present study, we examined how *env* sequence diversity of contemporary HIV-1 Indian clade C (isolated between 2020 and 2023) differentiates them from historical viruses as well as contemporary HIV-1 clade C of South African origin. To encompass contemporary HIV-1 of Indian origin at the population level, we obtained samples as source of HIV from nine geographically distinct origins representing different risk group. Although region-specific numbers of viruses were moderate, perhaps accounting for the fact that we saw no region-specific clustering, to the best of our knowledge, this is first such study of genetic and neutralization profiles of contemporary viruses from geographically distinct regions in India. A larger sample size of region-specific circulating forms would provide more precise phylogenetic details.

Although the Indian contemporary clade C *envs* continue to cluster with historical viruses, we found a significant drift in the degree of their sensitivity to CAP256-VRC26.25 and PGDM1400, the two clinically relevant bnAbs that target V1/V2 apex region of the viral Env protein. Over 45% and 40% of the contemporary viruses were found to be resistant to CAP256-VRC26.25 and PGDM1400 respectively. Our observation is consistent with our earlier study [10] and that of South African clade C viruses [9]. Conversely, the contemporary India viruses showed increased sensitivity to PGT121 which is in contrast to previous observation in South African clade C viruses [9]. These differences could be due to increased predicted glycosylation in *gp120,* particularly in V1/V2, as previously described [9, 22, 23] and possibly also differences in net V1 charges as observed in our study. The resistance to CAP256-VRC26.25 and PGDM1400 is also likely due to enrichment of resistance associated amino acid residues in the key contact sites on viral envelope protein; such enrichment of K166, Q169 and/or K169 residues in CAP256-VRC26.25 resistant viruses and D160, Q169 and K275 in PGDM1400 resistant viruses.

Mkhize *et al*. [9] recently also reported a trend in decreasing sensitivity of Africa clade C (obtained from the placebo arm of the AMP trial participants) to VRC01 and VRC07, an observation that was not noted with Indian clade C viruses tested in this study. These observations along with other *env* sequence features such as loop length, glycosylation and net charges that differentiated contemporary India and Africa HIV-1 clade C clearly indicates that they continue to evolve independently and distinctly at population level.

A notable observation made was that the second generation CD4bs bnAbs (N6, 1-18 and VRC07) were able to neutralize contemporary Indian clade C viruses with significantly better breadth and potency compared to the first generation CD4bs bnAbs (VRC01 and 3BNC117). They also were found to neutralize majority of the contemporary viruses that showed resistance to the V1/V2 apex directed bnAbs (CAP256-VRC26.25 and PGDM1400) and VRC01 and 3BNC117. Such observation indicate that in comparison to V1/V2 directed bnAbs, the key contact sites and epitopes for N6, 1-18 and VRC07 are evolutionarily preserved. The poor neutralization breadth conferred by VRC01 and 3BNC117 could possibly be because of the substitutions of amino acid residues resulting due to selection pressure during the course of natural infection at one or more of their key contact sites that were reported to be essential for their ability to neutralize efficiently [24–26]. Interestingly, the second generation CD4bs bnAbs (N6, 1-18 and VRC07) were found to neutralize the VRC01 and 3BNC117 resistant viruses with over 2-fold lower potency than what was observed with their corresponding sensitive viruses. This could possibly due the following reasons observed with few VRC07, N6 and 1-18 resistant viruses: (a) increased net charge in V1/V2 hypervariable regions (VRC07), differences in PNGS content in V1/V2 region and increased V1/V2 net charge in V/1V2 (N6) and/or (b) enrichment of resistance associated residues.

Combination of best-in-class bnAbs with distinct specificities have been reported to improve the optimal neutralization coverage of the HIV-1 diversity both by prediction and real-world application in experimental trials [27–31]. Emergence of HIV-1 clade C variants that showed broad resistance to major clinically relevant bnAbs was an interesting observation to note. Although few identified in this study, their presence in early infected individuals may imply that such resistant viruses can transmit and establish infection. Moreover, more such broadly resistant viruses are likely to evolve over time at the population level. It is therefore important to identify bnAbs that can be included in the antibody cocktail that can suitably compensate the inability of the existing best-in-class bnAbs to neutralize such viruses. Identification of viruses that are broadly resistant to existing best-in-class bnAbs also provides an opportunity to isolate new class of antibodies with new target specificities that are capable of neutralization evolving viruses that are broadly resistant to the existing bnAbs. Although we have predicted, based on individual virus neutralization data that BG18+N6+PGDM1400 would provide maximal coverage, however this needs to validated by neutralization assays.

One of the interesting observations made in this study is identification of mutations in *pol* gene associated with drug resistance in isolates obtained from over 10% ART naïve donors (**Table S1**). This indicates the ability of establishment of infection by drug resistant viruses [32, 33] which can potentially minimize the efficacy of antiretroviral therapy post exposure. Such observation further justifies the importance of the using next generation bnAbs as prevention strategy. One of the limitations of this study is that we examined neutralization properties of the cross-sectionally collected HIV+ samples and it will be useful to monitor longitudinally followed up cohort to periodically assess how HIV evolution overtime influences effectivity if the clinically relevant bnAbs. Moreover, several bnAbs that are under clinical development were isolated long ago and emerging reports indicate including our present study indicates several of them becoming irrelevant for intervention. Therefore, the need for periodic assessment of sequence and neutralization profiles of the regionally relevant contemporary HIV-1 forms against engineered optimized clinically relevant bnAbs is required for prioritizing and development of effective bnAbs as product for prevention.

## Materials and Methods

### Ethics statement

All clinical samples from nine different sites in India were obtained following approval from respective institutional ethical committees. Written informed consent forms in English and local languages were provided and duly signed by all the recruited study participants. Experiments at respective institutions were initiated post approval of institutional ethics committee. All experiments were carried out at the THSTI, Faridabad post approval of institutional ethics committee (IEC) and institutional biosafety committee.

### Study participants

HIV-1 infected individuals were recruited from nine geographically distinct clinical sites in India. They are from Eastern (Kolkata), Western (Mumbai, Belagavi), Northern (Delhi, Bhopal), Southern (Nellore, Hyderabad, Vizag) and North Eastern (Aizawl). following approvals from the institutional ethics committee at all participating institutions. A total of 232 study participants were recruited from nine different geographical sites as indicated in **Table 1**. Clinical parameter data such as CD4 counts, viral load and antiretroviral therapy status were obtained for each study participant.

### Plasmids, antibodies and cells

Plasmids encoding full length codon optimized *gp160* of Indian origin synthesized at GenScript Inc. were used for preparing pseudoviruses. Plasmids encoding HIV-1 clade C *env* genes of South African origin from AMP placebo arm reported earlier [9] were used to prepare pseudoviruses for the neutralization assay. pSG3ΔEnv was obtained from the NIH AIDS Reagent and Reference Program. Plasmids encoding heavy and light chain immunoglobulins of CAP256-VRC26.25 was provided by Prof Lynn Morris and ones with VRC01, VRC07, N6, 1-18, PGDM1400, 3BNC117, BG18, 10-1074, 10E8, VRC34.1 were provided by the IAVI Neutralizing Antibody Center. HEK 293T, TZM-bl, were obtained from the American Type Culture Collection (ATPC) and GHOST-Hi5, GHOST-CXCR4 cells were obtained fromn the NIH AIDS Reagents & Reference Program respectively. GHOST-CCR8 cells were kindly provided by Paul Clapham. Expi293 cells were purchased from Thermo Inc.

### Isolation of viral and genomic DNA and cDNA synthesis

Viral RNA was isolated from plasma using the High Pure viral RNA kit (Roche) as per manufacturer’s instruction as described earlier[34]. Genomic DNA was isolated from peripheral blood mononuclear cells (PBMC) using QIAmp blood DNA mini kit (Qiagen) as per the manufacturer’s instructions and as described earlier [34]. Plasma isolated RNA was primed with EnvR1 oligo (5’- GCACTCAAGGCAAGCTTTATTGAGGCT-3’) proximal to 3’ end of the HIV RNA genome (HXB2: 9605-9632) and Aenvseq4 (5’- CAAGCTTGTGTAATGGCTGAGG -3’) binding downstream of the *pol* gene (HXB2: 6817-6838). Synthesis of cDNA was performed using Superscript III first strand synthesis kit (Invitrogen) following the protocol provided by the manufacturer.

### Amplification of full length *gp160 and pol*

Full length *env* (*gp160*) genes were PCR amplified from HIV + plasma samples with slight modification as described previously [34]. *Rev-env gp160* cassette were amplified from the cDNA product using La Taq high fidelity DNA polymerase in the 1st round (Takara Bio Inc.) and PrimeSTAR GXL high fidelity DNA polymerase (Takara Bio Inc.) in the second round. The primers used for the 1^st^ round were EnvF1: 5′- AGARGAYAGATGGAACAAGCCCCAG-3′ (HXB2: 5550–5574) and EnvRP2: 5′-GTGTGTAGTTCTGCCAATCAGGGAA -3′ (HXB2: 9157–9181) while for the second round were Env IF: 5′-CACCGGCTTAGGCATCTCCTATGGCAGGAAGAA -3′ (HXB2: 5950– 5982) and EnvIR: 5′-TATCGGTACCAGTCTTGAGACGCTGCTCCTACTC -3′ (HXB2: 8882–8915). PCR condition followed for both rounds were: initial denaturation of 94 °C for 2 min followed by 15 cycles of 94 °C for10 secs, 60 °C for 30 secs, 68 °C for 3 min, 20 cycles of 94 °C for 10 secs, 55 °C for 30 s, 68 °C for 3 min with final extension of 68 °C for 10 min. The *gp160* amplicons were purified and subsequently subjected to short (Illumina) and short read deep sequencing to obtain dominant sequences as described below which were subjected to codon optimization, synthesized and cloned into pcDNA3.1 expression vector. Few *env* clones (**Table 1**) were cloned in-house in pcDNA3.1/V5-His-TOPO (Invitrogen Inc.) vector as described before [34]. The primers used for the 1^st^ round towards pol amplification were Pro5F: 5′- AGAAATTGCAGGGCCCCTAGGAA -3′ (HXB2: 1996-2018) and PolR1: 5′- GGTACCCCATAATAGACTGTRACCCACAA -3′ (HXB2: 6324-6352) while for the second round were Pro3F: 5′- AGANCAGAGCCAACAGCCCCACCA -3′ (HXB2: 2143-2166) and PolR2: 5′- CTCTCATTGCCACTGTCTTCTGCTC -3′ (HXB2: 6207-6231). PCR condition followed in the both rounds were initial denaturation of 94 °C for 2 min followed by 15 cycles of 94 °C for10 secs, 65 °C for 30 secs, 68 °C for 3 min, 20 cycles of 94 °C for 10 secs, 55 °C for 30 s, 68 °C for 3 min with final extension of 68 °C for 10 min.

### Next generation deep sequencing and construction of *env* sequences

Env amplicons were sequenced using both long read Oxford Nanopore (ON) and short read Illumina (IL) platforms. Next generation sequencing was performed for 5’ fragments using the Illumina platform while 3’ fragments were sequenced using both Illumina and Oxford nanopore platforms. The raw data obtained from the nanopore sequencing was converted to Fastq files using Guppy basecaller (v6.3.7). Raw reads were further filtered for quality and read length using Prowler (Flags: -l 1500 -q 12 -c “LT” -g “F1” -m “S”) [35]. The reads were aligned to the HIV-1 subtype C reference sequence (GenBank ID: AF067155.1) [36] using Minimap2 [37, 38] and processed for read sorting and filtration with samtools [39]. Reads encompassing the entire gene were extracted from the binary alignment maps using Picard tools (https://broadinstitute.github.io/picard/). Reads were further clustered and corrected using isONclust and isONcorrect respectively into quasispecies clusters [40, 41]. Consensus sequences were generated for each quasispecies clusters using iVar [42]. Quasispecies thus constructed were further corrected with the help of Illumina reads using Pilon [43].

### Preparation of Env-pseudoviruses

Pseudotyped viruses were prepared as described previously [44]. Briefly 293T cells were co-transfected by plasmid DNA encoding *gp160* and pSG3ΔEnv plasmid (having a premature stop codon at the beginning of *env*) into 293T cells in 6-well tissue culture plates using FuGENE6 transfection reagent kit (Promega Inc.). Cell culture supernatants containing pseudotyped viruses were harvested at 48 h post transfection and subsequently stored at -80° C until use. The virus infectivity was measured using TZM-bl reporter cells by addition of pseudoviruses containing DEAE-dextran (25 µg/ml) in 96-well microtiter plates, and the viral titers were determined by measuring the luciferase activity using Britelite luciferase substrate (PerkinElmer Inc.) with a Victor X2 luminometer (PerkinElmer Inc.).

### Coreceptor usage

Coreceptor preference of contemporary envelops was examined by cell-cell fusion assay as described before [45]. Briefly, 293T cells expressing individual *env* was mixed with GHOST-Hi5, GHOST-CXCR4 and GHOST-CCR8 post 24 hours of transfection and further incubated for additional day at 37°C in a CO_2_ incubator. Syncytia forming giant cells were identified by staining with chilled methanol containing 1% methylene blue and 0.25% basic fuchsin. 293T cells expressing 16055-2.3 for GHOST-Hi5 [46], NARI-VB105 [45] for GHOST-CXCR4 and NARI-VB52 for GHOST-CCR8 [47] were used as positive controls.

### Pseudovirus Neutralization assay

Neutralization assays were carried out using TZM-bl cells as described before [44]. Briefly, Env-pseudotyped viruses were pre-incubated in 96-well tissue culture plates with various concentrations of bnAbs (IgG) for an hour at 37°C in a CO_2_ incubator under humidified conditions. Subsequently, 1 × 10^4^ TZM-bl cells were added to the mixture in the presence of 25 µg/ml DEAE-dextran (Sigma, Inc.). The plates were further incubated for 48 h. The degree of virus neutralization was assessed by measuring reduction in relative luminescence units (RLU) in a luminometer (Victor X2; PerkinElmer Inc.). The IC50 and IC80 values were calculated using R using the DRC statistical package (analysis of dose response curves.

### ARV resistance mutations prediction

Illumina FASTQ reads were filtered for quality (>Q30) using Trimmomatic (v0.39). All the reads were aligned to the HXB2 genome using bwa-mem (v0.7.17-r1188). BAM files were filtered for quality using samtools. Variant calling was performed for the *pol* gene using iVar pipeline. Drug resistance mutation prediction was then performed for the variants obtained using Stanford drug resistance database HIVdB (https://hivdb.stanford.edu/hivdb/by-patterns/). Resistance patterns were recorded only for variants with frequency greater than 10%.

### Phylogenetic analysis

Phylogenetic trees were generated for 249 HIV-1 envelope amino acid sequences that included 232 contemporary sequences from India and 17 HIV-1 group M subtype reference sequences; and 594 HIV-1 envelope amino acid sequences that included 233 contemporary and 132 historical sequences from India, 74 contemporary and 138 historical sequences from Africa along with 17 HIV-1 group M subtype reference sequences. These sequence datasets were aligned using MAFFT and the alignment was manually curated in BioEdit v7.2.5. The tree was constructed with IQ-TREE under HIVb model [48, 49] with estimated Ƴ parameters and number of invariable sites. The robustness of the tree topology was further assessed by SH-aLRT as well as 1000 ultrafast bootstrap replicates implemented in IQ-TREE as described earlier [17].

### Variable region characteristics and prediction of pNLG

Variable region characteristics such as loop length, charge and number of pNLG sites were assessed for all envelope sequences using the ‘variable characteristics tool’ hosted at the Los Alamos National Laboratory HIV database (LANL-HIVDB, https://www.hiv.lanl.gov/content/sequence/VAR_REG_CHAR/index.html). Potential N linked glycosylation sites prediction was performed with the tool N-Glycosite at LANL-HIVDB (https://www.hiv.lanl.gov/content/sequence/GLYCOSITE/glycosite.html).

### bnAb contact site assessment

For bnAbs CAP256-VRC26.25, PGDM1400, PGT145 (V2 apex directed), PGT121, BG18, 10-1074

(V3g supersite directed), VRC01, VRC07, 1-18, N6, 3BNC117 (CD4 binding site) and 10E8 (MPER directed), specific epitope contact positions as well as documented sensitivity/resistance imparting variants at each position were retrieved from CATNAP database (https://www.hiv.lanl.gov/components/sequence/HIV/neutralization/main.comp). Each of the sequences were then assessed for presence of sensitive/resistant/undefined mutation at each of these positions using custom bash scripts. In sequence logos, O has been used to differentiate potential N linked glycosylated Asparagine from potentially unglycosylated Asparagine (N).

### CombiNAber analysis

Optimal combination prediction was performed with the CombiNAber tool at LABL-HIVDB (https://www.hiv.lanl.gov/content/sequence/COMBINABER/combinaber.html). CombiNAber predictions were made with the IC50 and IC80 neutralization data using the bliss hill model at target concentrations of 10ug/mL and 1ug/mL for 3 distinct specificity bnAb combinations as well as active coverage by at least 2 bnAbs.

### Statistical analyses and data presentation

Phylogenetic trees were annotated using the ‘ggtree’ package in R. Sequence logos were constructed with ‘ggseqlogo’ package in R. All plots were prepared using the R package ggplot2. Statistical comparison of variable region characteristics with Mann-Whitney test. Fisher’s test for abundance of bnAb resistance associated residues was performed through R statistical computing software (v3.4.0) and R studio v1.0.143. Statistical analysis for neutralization breadth and potency were done using GraphPad Prism version 10 for Windows, GraphPad Software.

## Data availability

Novel *env* and *pol* nucleotide sequences obtained from Indian donors are being submitted to GenBank. *env* nucleotide sequences from FRESH cohort has been submitted to GenBank.

## Individual contributions

JB with help of VP, KGM conceptualized the study and supervised the entire study, was responsible for funding acquisition and resources for the conduct of the present study; JS, PJ, RM along with JB planned major experiments; JS, PJ, RM carried out majority of the experiments and carried out sequence and neutralization data analysis; JS led the next generation sequencing based experiments along with SB, PK, SS, NP, constructed consensus sequences used for synthesis of constructs used for preparation of pseudoviruses; NN, VP, PD, NK, SB, VB; MA, CVK, TRD helped with viral gene amplification and sequencing; PJ, JS, RM, SD carried out molecular cloning; SC, ST, BS, SD helped with preparation of pseudoviruses and neutralization assays; SM, NS, CP, NK prepared purified IgGs and carried out QA/QC for ensuring quality and bnAb specificities before they used in neutralization assays; AKS, JS, SM, BB, SKG contributed in creation of cohorts in different sites in India and supervised sample preparation, clinical investigations; DK, DB, DP, RSP, SM, H, RD, AK helped with separation of plasma and PBMCs, assessment of immunological parameters; DKit, HK, NNM carried out majority of neutralization assays and sequence analysis of HIV-1 clade C viruses of South African origin; BN and KG helped with providing HIV-1 clade C *env* sequence information obtained from FRESH acute cohort, South Africa; PLM and TN helped with data analysis, supervision of experiments carried out at individual laboratories in South Africa; DS helped with experimental design and data analysis; JB with assistance from JS, PJ, RM, VP, KGM, PLM and TN wrote the manuscript.

## Acknowledgments.

First, we sincerely thank the study participants for consenting to provide clinical materials for this study. We gratefully acknowledge the support rendered by everyone at all the nine sites from where clinical materials were obtained and the members of all participating laboratories for support with experiments. We thank Paramita Saha, Joyeeta Mukherjee, Monal Nagrath, Shweta Chatrath, Elise Landais, Tanvi Khera, Rajat Goyal, S Saravanan, Raghavan Sampathkumar and Sai Shankar Ramakrishnan for operational supports. We thank IAVI Neutralizing Antibody Center for providing reagents for carrying our experiments. Special thanks to the National AIDS Control Organization (NACO), Ministry of Health & Family Welfare, Govt. of India for their support to include NACO-ART centers for collection of clinical samples for this particular study.

## Supporting data

### Supporting Tables

**Table S1.** Demographics details of nine different geographical regions in India, risk groups and ART status of HIV+ donors.

**Table S2.** Neutralization breadth and potency (IC50) of pseudoviruses expressing contemporary India clade C *envs*.

### Supporting Figures

**Fig.S1.** Hierarchal clustering using heatmap depicting the magnitude (https://www.hiv.lanl.gov/content/sequence/HEATMAP/heatmap.html) of neutralization sensitivity of contemporary HIV-1 India clade C viruses against 14 bnAbs with distinct epitope specificities. Heatmap was prepared using IC50 and IC80 values (ug/mL).

**Fig.S2.** Comparison of Env variable loop, PNGS and net charges between South Africa clade C viruses that showed sensitivity and resistance to CAP256-VRC26.25 and PGDM1400.

**Fig.S3.** Sequence features of India clade C viruses encoding contemporary *envs* sensitive and resistant to V3 glycan supersite directed bnAbs. A. Sequence logos of PGT121, 10-1074 and BG18 sensitive and resistant viruses. B. Variable loop, PNGs and net charge characteristics between sensitive and resistant viruses.

**Fig.S4.** Sequence logo of India clade C viruses encoding contemporary *envs* sensitive and resistant to the CD4bs directed bnAbs.

**Fig.S5.** Comparison of variable loop length, PNLGs and net charge of contemporary Indian clade C viruses sensitive and resistant to CD4bs directed bnAbs.

**Fig.S6.** Comparison of variable loop length, PNLGs and net charge between Indian and South Africa clade C viruses resistant to CAP256-VRC26.25 and PGDM1400.

**Fig.S7.** Comparison of frequency of key contact residues on *envs* that when expressed as pseudoviruses of India (N=115) and South Africa (N=40) origins showed resistance to V3 glycan directed bnAbs.

**Fig.S8.** Abundance of CAP256-VRC26.25/PGDM1400 sensitivity/resistance associated residues in sequences from India (N=118) and South Africa (N=41) that have not been tested through *in vitro* neutralization assays. X axes define amino acid positions in gp160 whereas Y axes define percent abundance of residues. The tables indicate the residues associated with sensitivity or resistance. Und: undefined.

